# rdrugtrajectory: An R Package for the Analysis of Drug Prescriptions in Electronic Health Care Records

**DOI:** 10.1101/2021.01.08.425952

**Authors:** Anthony Nash, Tingyee E. Chang, Benjamin Wan, M. Zameel Cader

## Abstract

Primary care electronic health care records are rich with patient and clinical information. Studying electronic health care records has resulted in marked improvements to national health care processes and patient-care decision making, and is a powerful supplementary source of data for drug discovery effort. We present the R package **rdrugtrajectory**, designed to yield demographic and patient-level characteristics of drug prescriptions in the UK Clinical Practice Research Datalink dataset. The package operates over Clinical Practice Research Datalink Gold clinical, referral and therapy datasets and includes features such as first drug prescriptions analysis, cohort-wide prescription information, cumulative drug prescription events, the longitudinal trajectory of drug prescriptions, and a survival analysis timeline builder to identify risks related to drug prescription switching. The **rdrugtrajectory** package has been made freely available via the GitHub repository.

## 1. Introduction

The UK Clinical Practice Research Datalink (CPRD) service offers high quality longitudinal data on 50 million patients with up to 20 years of follow-up for 25% of those patients. The service provides drug treatment patterns, feasibility studies and health care resource use studies. Patient electronic health care records (EHR) are stored as coded and anonymised data and sourced from over 1,800 primary care practices across England. CPRD holds information on consultation events, medical diagnoses, symptoms, prescriptions, vaccination history, laboratory tests, and referrals. CPRD can provide routine linkage to other health-related patient datasets, for example: Small area level data, such as patient and/or practice postcode linked deprivation measures; data from NHS digital which includes hospital episode statistic, outpatient and accident and emergency data; and cancer data from Public Health England. Evidence from EHRs is making an impact on primary care decision-making and best practice Oyinlola *et al*. (2016). With nationwide longitudinal datasets more readily available, the evaluation of treatments over long timescales can contribute to clinical decision-making Hepp *et al*. (2017). For example, adverse events caused by prescription medication can be studied using retrospective data in situations where randomized clinical trials may prove impractical Ghosh *et al*. (2019); Bally *et al*. (2017).

This publication serves as an introduction to the **rdrugtrajectory** R package and whilst this publication is by no means a complete tutorial, we will expand on some of the main package features, such as, how to: Isolate patients by first drug prescriptions at given clinical events; calculate time-invariant prescriptions; construct survival analysis timelines (compatible with Cox proportional hazard regression and Kaplan Meier curves), and; visualise patient prescription switching. For a comprehensive list of functions please visit the Github repository https://github.com/acnash/rdrugtrajectory. Almost all features can be controlled by covariates or stratified by some variable, for example, by gender, age, medical codes or treatment product codes.

The example code, figures and data structures presented here mimic a small fraction of our own research. In the interest of patient confidentiality, the clinical data used in the analysis have been fabricated. We present a brief tour of some of the functions available, starting with a discussion on the CPRD data structure and how records must be formatted. A glossary of terms has been provided (Table 1) to assist the reader.

**Table 1:** Table of frequently used terms.

| Term | Description |
| --- | --- |
| **rdrugtrajectory** | An R packaged designed for the management of CPRD prescription data. |
| *clinical* | The *ClinicalNNN.txt* dataset presented in a **rdrugtrajectory** dataframe. |
| *referral* | The *ReferralNNN.txt* dataset presented in a **rdrugtrajectory** dataframe. |
| *therapy* | The *TherapyNNN.txt* dataset presented in a **rdrugtrajectory** dataframe. |
| *AdditionalNNN.txt* | The CPRD dataset of additional clinical information, for example, patient smoking status and alcohol consumption. Data can be retrieved using *CPRDLookups.R*. |
| *modecode* | A CPRD identifier that denotes medical conditions, diagnosis and complaints made by a patient. *medcodes* are recorded in the *ClinicalNNN.txt* and *ReferralNNN.txt* files. |
| *procode* | A CPRD identifier that denotes treatment products, including drugs, foods, and medical apparatus. *prodcodes* are recorded in the *TherapyNNN.txt* files. |
| *patid* | A unique CPRD patient identifier. Used to link datasets. |
| *event* | Any *procode* or *medcode* in a patient's EHR. |
| *eventdate* | The date of an event recorded by a general practitioner. Present in all three datasets and corresponding **rdrugtrajectory** dataframe. |
| *IMD* | Index of Multiple Deprivation score - a UK Government socioeconomic measurement based on postcode of the clinic or a patient's registered address. |
| Prescription | A general time for any *procode* prescribed for treatment. |
| *medical history* | Indicates a combination of one or more sets of CPRD data, for example, the collection of all clinical and therapy EHR for patients with a *medcode* for migraine. |
| *product.txt* | A plain text file that contains all *prodcodes* with a description and comes bundled with the CPRD Data Dictionary. The file is used to link a *procode* with a description. |

## 2. **rdrugtrajectory** package and data structures

### 2.1. rdrugtrajectory availability and installation

**rdrugtrajectory** is free to download from the Github repository https://github.com/acnash/rdrugtrajectory and holds an MIT license. Fabricated CPRD clinical and CPRD prescription records in addition to age, gender and index of multiple deprivation scores are included for test and tutorial purposes. Before installing the package, the following R dependencies are required: `plyr`, `dplyr`, `foreach`, `doParallel`, `data.table`, `parallel`, `splus2R`, `rlist`, `reda`, `ggplot2`, `ggalluvial`, `stats`, `utils` and `useful`. The latest **rdrugtrajectory** binary is install using:

~~~
install.packages(“path/to/tar/file”, source = TRUE, repos=NULL)
~~~

**rdrugtrajectory** was developed and tested on R version 4.0.1. Please consult the Github page for release notes, the latest version and up to date installation instructions.

### 2.2. CPRD product descirption

Several **rdrugtrajectory** functions use the CPRD *product*.*txt* file for assigning a text description to a prescription *prodcode*. The *product*.*txt* (and *medical*.*txt* for *medcode* description) is available in the CPRD Data Dictionary Windows software. It is important that the file remains in plain text, with columns tab-delimited. The files can be simplified by removing all non-essential products. Finally, all the eleven columns that make up the *product*.*txt* file must be available, with the first column containing all *prodcodes* and the fourth column containing the *product description*. A simplified *product*.*txt* file, presented below, can be downloaded from the Github page.

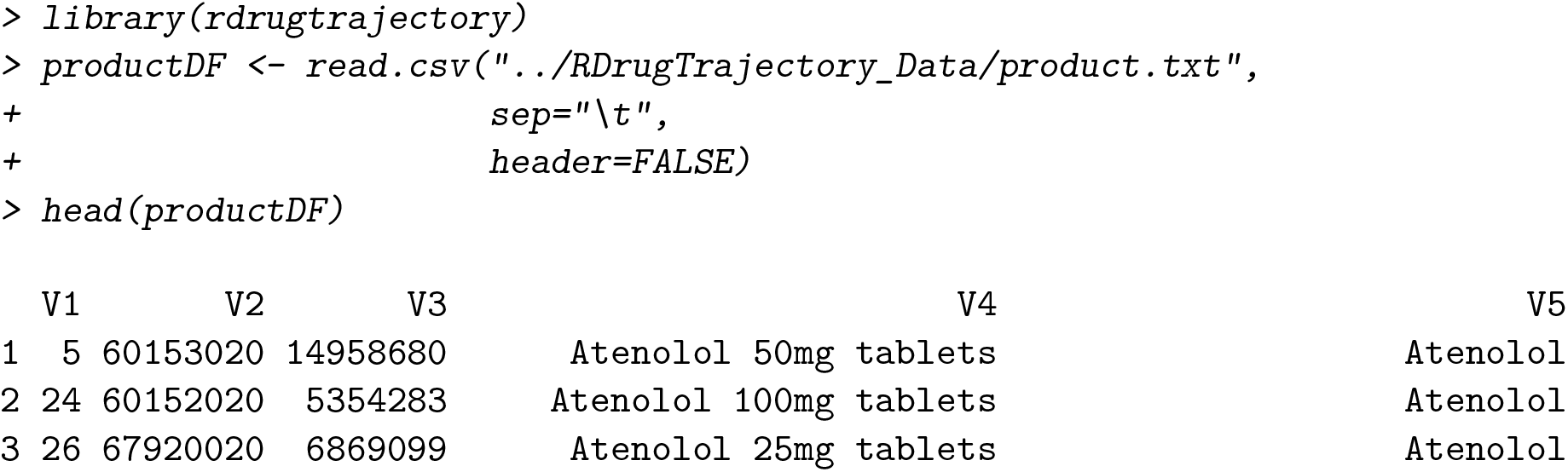
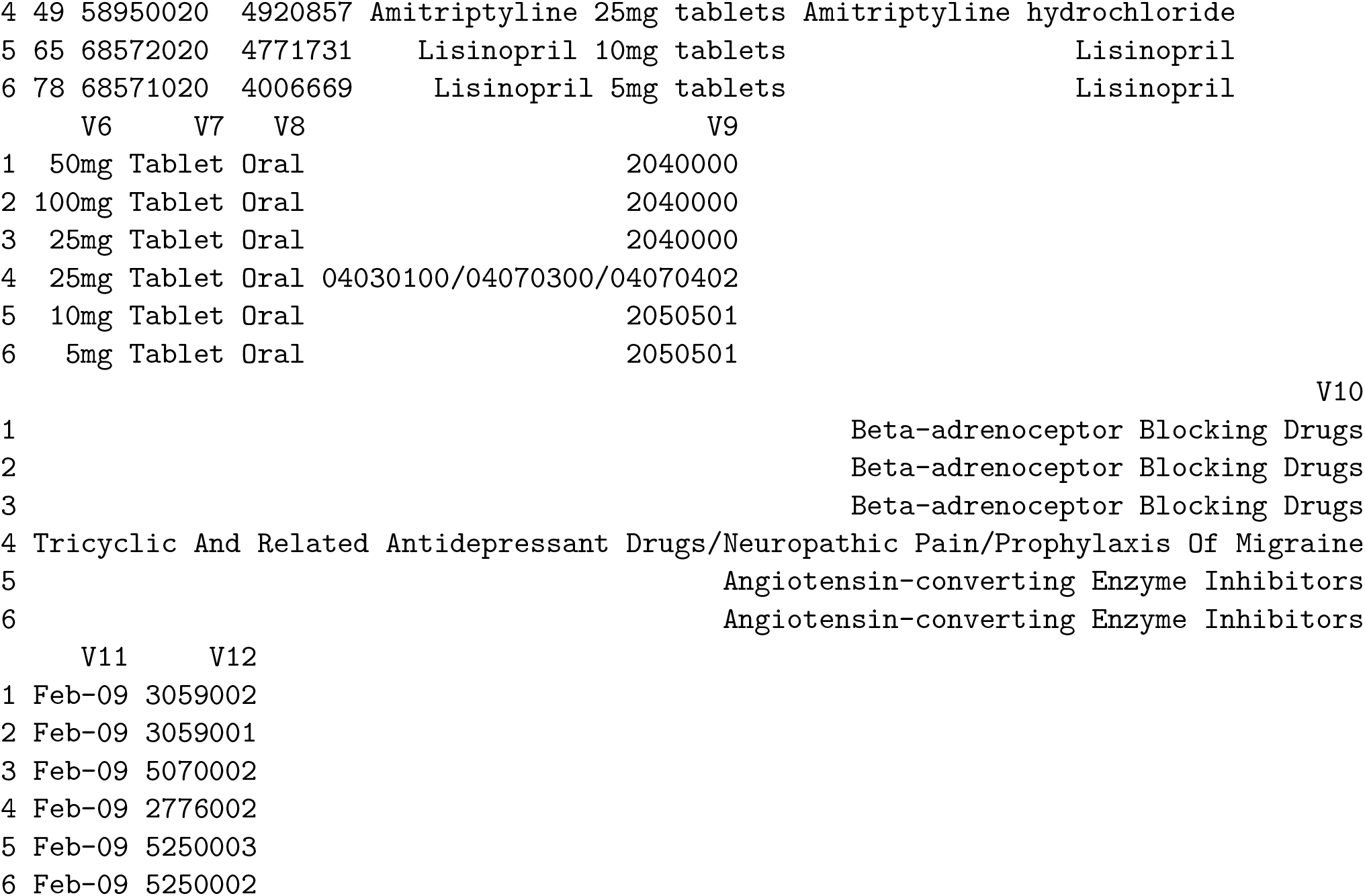

**Figure 1:**
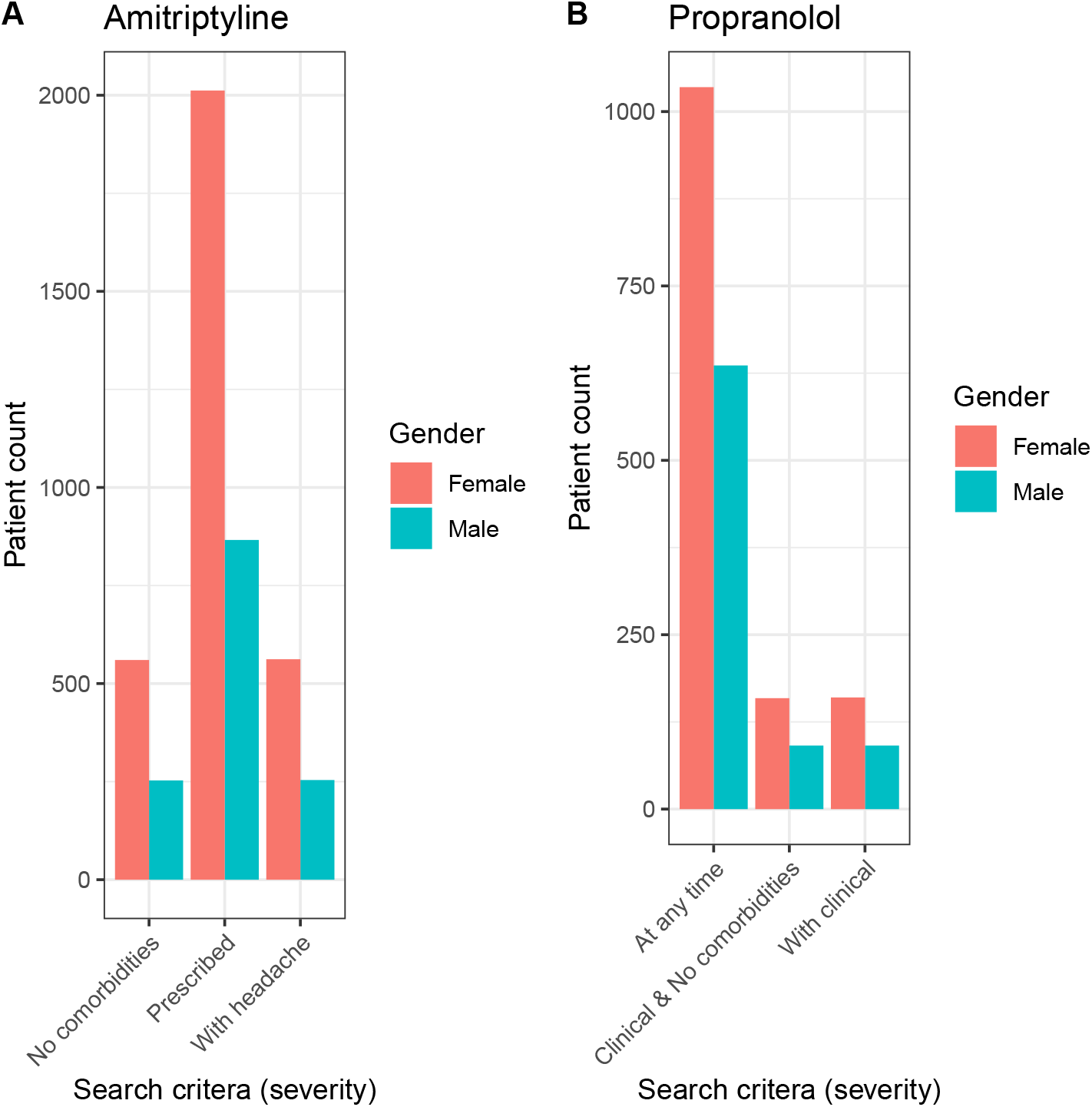
The number of patients prescribed (A) amitriptyline or (B) propranolol. The criteria to match against clinical data is indicated: at any time, with a clinical record, and with a clinical record clear off topic clinical events.

**Figure 2:**
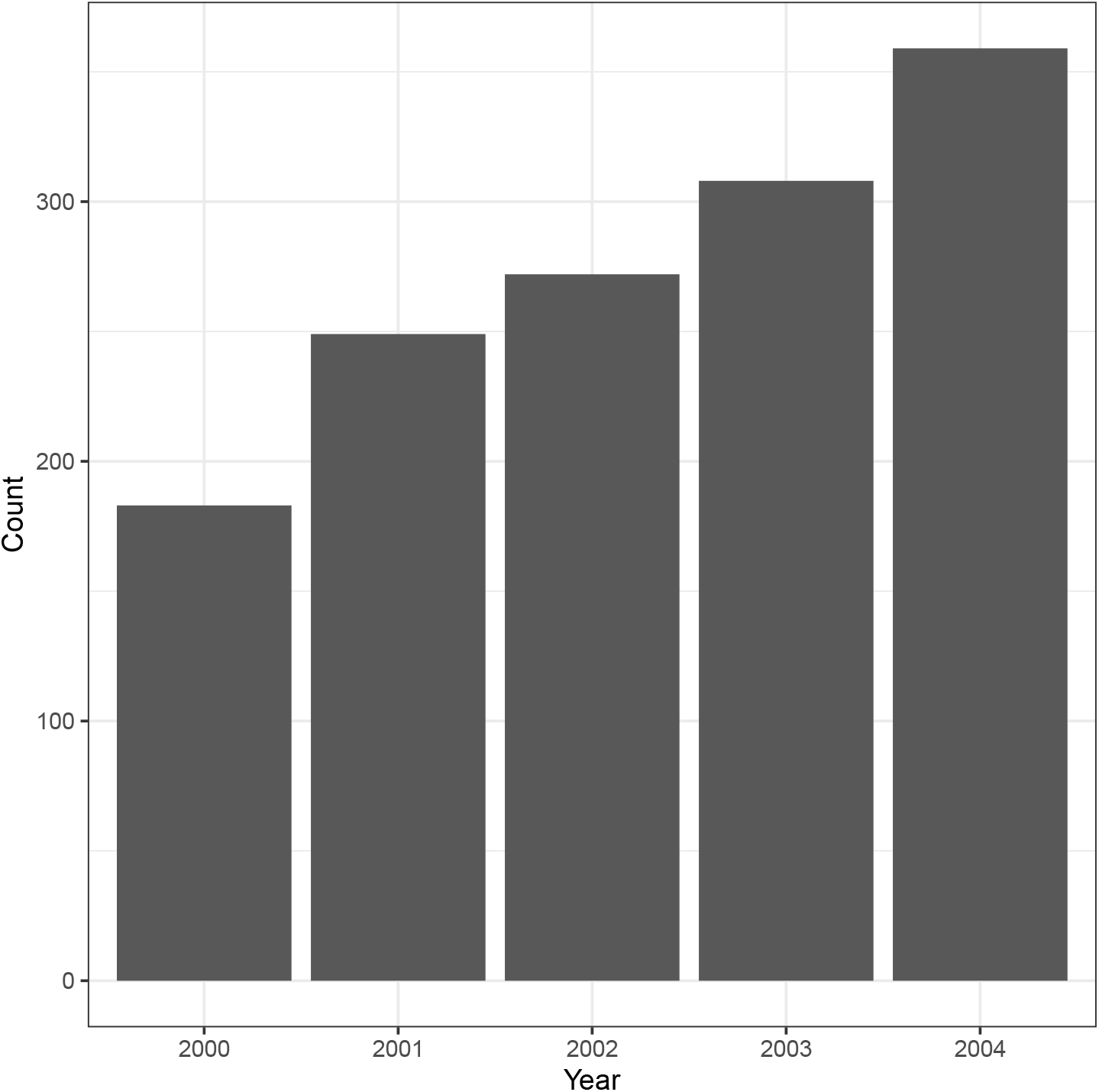
The number of patients prescribed amitriptyline from the start of the year 2000 to the end of 2004, stratified in year intervals.

### 2.3. rdrugtrajectory package structure

**rdrugtrajectory** contains three R files: (1) all functions related to data curating and searching reside within *PRDDrugTrajectory*.*R*; (2) analysis tools and timeline construction reside within *CPRDDrugTrajectoryStats*.*R*; and, (3) all utilities including input/output operations reside within *CPRDDrugTrajectoryUtils*.*R*. The packages contains several fabricated CPRD datasets: `testClinicalDF`, `testTherapyDF`, `ageGenderDF`, `imdDF`, and `drugListDF`. A description of each, along with information on data types and structures are given below.

### 2.4. The CPRD EHR data structure

The structure of CPRD Gold data may depend on whether the CPRD license holder performs intermediate data management steps before releasing data to the user. However, typically, CPRD Gold data follows the CPRD Gold specification https://cprdcw.cprd.com/_docs/CPRD_GOLD_Full_Data_Specification_v2.0.pdf. Currently, **rdrugtrajectory** supports EHR data from the flat files *ClinicalNNN*.*txt, ReferralNNN*.*txt*, and *TherapyNNN*.*txt*. The Additional Clinical Details files (*AdditionalNNN*.*txt*) are currently supported using our released R script *CPRDLookups*.*R* https://github.com/acnash/CPRD_Additional_Clinical **?**. Patients are assigned a unique numerical *patid* value. The operations performed by **rdrugtrajectory** requires the *patid* to identify patients and subset patient groups. We recommend that *patid, medcode, prodcode* are kept as character data throughout any preliminary data curating steps. Medical events are recorded as codes and stored in the *ClinicalNNN*.*txt* and *ReferralNNN*.*txt* under the column header *medcode*. Prescription events, such as drug prescriptions are also recorded as codes and stored in the *TherapyNNN*.*txt* file under the column header *prodcode* and the sequences of repeat prescriptions are under the *issueseq* column header. Dates associated medical and prescription events, recorded by the General Practitioner, are stored under the column header *eventdate*.

### 2.5. Essential data types and data structures

**rdrugtrajectory** can operate over CPRD Gold EHR clinical, referral and prescription data provided each dataset format is presented as separate R dataframes or combined into a **rdrugtrajectory** *medical history* dataframe. The construction of clinical, referral and prescription dataframes require, as a minimum, a *patid* and *eventdate* column, and either *medcode* or *prodcode* (for therapy data, *issueseq* is necessary), and presented in that order. Every record of *medcode* or *prodcode* must be accompanied by an *eventdate* entry (encoded as a `Date` class of the form *YYYY-MM-DD*). Patients can have duplicate events within the same data set and between data sets. Medical and prescription codes can be retrieved from the corresponding *medical*.*txt* and *product*.*txt* files which come bundled with the CPRD Data Dictionary Windows application. **rdrugtrajectory** comes packaged with fabricated EHR data in the structure of:

~~~
*> library(rdrugtrajectory)
> #fabricated clinical data (referral data follows the same format)
> names(testClinicalDF)*
[1] “patid” “eventdate” “medcode” “consid”
*> #fabricated prescription data
> names(testTherapyDF)*
[1] “patid” “eventdate” “prodcode” “consid” “issueseq”
~~~

Users can check if the structure of an EHR dataframe meets the requirements for this package by calling `checkCPRDRecord`; additional columns such as consultation identification number (*consid*) are not considered. In the following instance, a prescription dataset with the required columns and the optional consultation identification number is presented.

~~~
*> library(rdrugtrajectory)
> #check the structure of testTherapy, specify that it is therapy data
> checkCPRDRecord(df=testTherapyDF, dataType="therapy”)*
[1] “The data.frame is appropriately formatted. Returning TRUE.”
[1] TRUE
*> #display the rdrugtrajectory EHR therapy dataframe
> str(testTherapyDF, strict.width=“wrap”)*
‘data.frame’: 91647 obs. of 5 variables:
$ patid : int 3515 3515 3515 3515 3515 3515 3515 3653 3653 3653 ...
$ eventdate: Date, format: "2005-02-24" "2006-01-26" ...
$ prodcode : int 83 83 83 707 707 707 707 297 297 297 ...
$ consid : int 540850 540865 540892 541108 541114 541118 541133 571336 571345
571357 ...
$ issueseq : int 0 0 0 0 0 0 0 0 1 2 ...
~~~

Users can combine with the **rdrugtrajectory** EHR dataframes any number of patient and EHR data to act as covariates and stratifying variables, typically this can be done using the R `cbind` operation. For example, BMI and smoking status, both of which can be retrieved from the *AdditionalNNN*.*txt* dataset files using *CPRDLookups*.*R*, can be linked by searching for and binding with the record *patid* values. The **rdrugtrajectory** package contains several utility functions to retrieve CPRD data, including, patient year of birth, gender (male or female) and either patient-level or clinical-level index of multiple deprivation score (IMD). The patient age can be determined by adding 1800 to the value in *yob* column in the *Patient* CPRD EHR dataset and then subtracting that value (birth year) from the year of the CPRD database release. This data requires preliminary treatment before presenting to the **rdrugtrajectory** package. Patient age, gender and IMD score must be presented in a dataframe with the linked patient column *patid*, along with the columns *age, gender*, and *score*. Providing the *patid* column is preserved, patient characteristics can be presented in separate dataframe, for example:

~~~
*> library(rdrugtrajectory)
> #patient age and gender as one dataframe
> str(ageGenderDF, strict.width=“wrap”)*
‘data.frame’: 3838 obs. of 3 variables:
$ patid : int 1 2 3 4 5 6 7 8 9 10 ...
$ yob : num 45 35 33 42 63 57 34 51 51 22 ...
$ gender: int 2 2 1 2 2 1 2 2 2 1 ...
*> #clinic-level IMD score as one datafrmae
> str(imdDF, strict.width=“wrap”)*
‘data.frame’: 2126 obs. of 3 variables:
$ patid : int 6 11 16 34 42 44 54 60 63 79 ...
$ pracid: int 184 31 66 344 66 47 18 90 379 317 ...
$ score : int 1 3 1 4 1 2 1 5 1 2 ...
~~~

The *patid* patient identifier is fundamental in every operation performed by **rdrugtrajectory**. The examples presented here and those in the reference manual rely on searching and subsetting EHR data using a list or vector of patient identifier. The function `getUniquePatidList` will retrieve an R `List` of patient identification numbers from any dataframe with a *patid* column.

The aforementioned **rdrugtrajectory** EHR dataframes, clinical, referral and therapy, can be combined into a single dataframe. We refer to this dataset instance as the patient’s *medical history* and can be constructed using `constructMedicalHistory`. This dataframe expects events to be in chronological order, and will introduce a new column, *code* and *codetype* to denote each of the combined events. The code (*medcode* and/or *prodcode*) can be distinguished by a *codetype* value of *c* (clinical events), *r* (referral events), and *t* (prescription events). Events are returned in chronological order using the *eventdate* data. The following code demonstrates how to retrieve a list of patient identifier from a prescription dataframe and from a *medical history* dataframe, followed by how to subset using base R operations and, finally, the *medical history* dataframe structure.

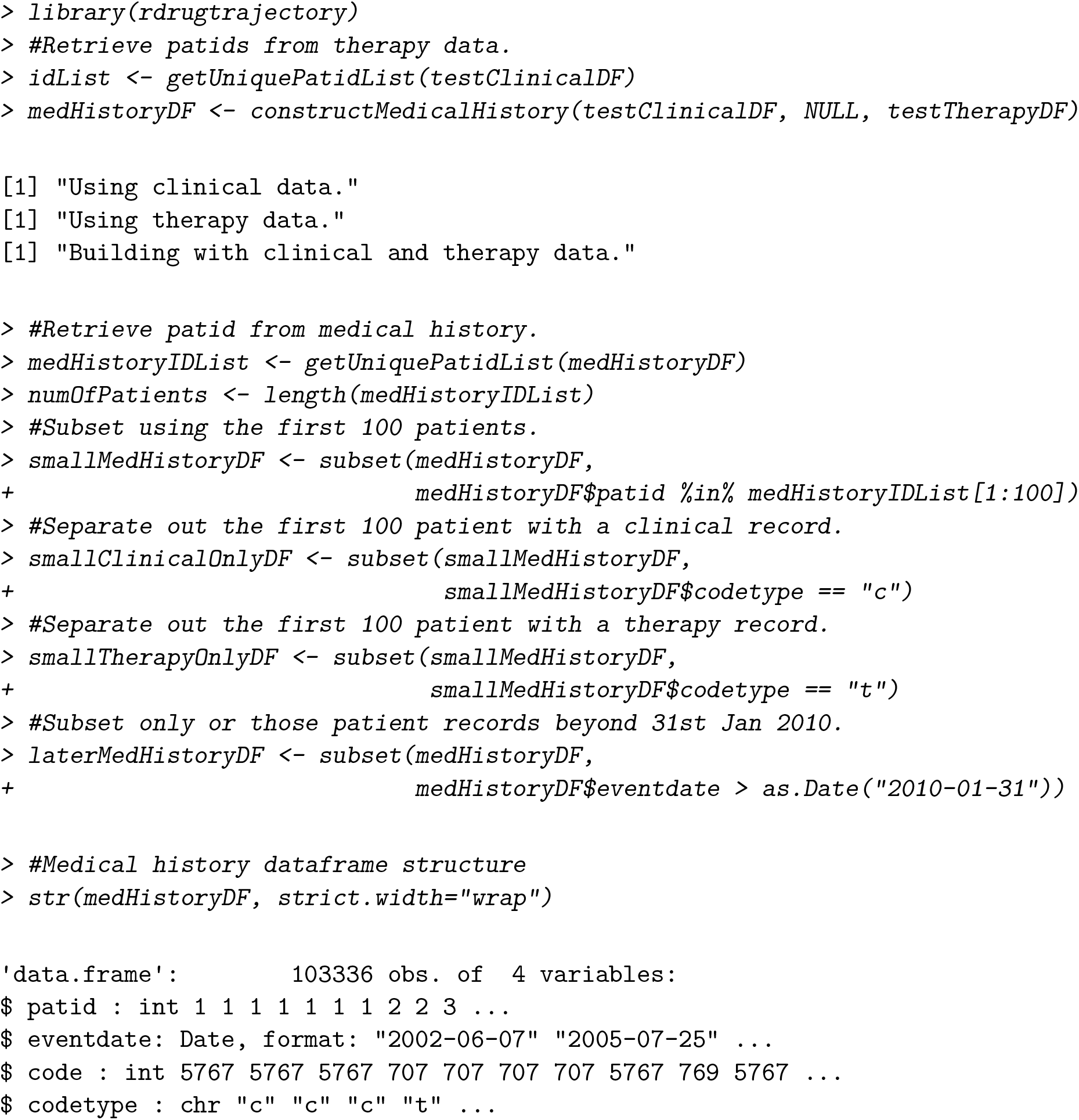

The *patid* data can also be used to retrieve patient characteristics, for example, the gender of the patient using `getGenderOfPatients`:

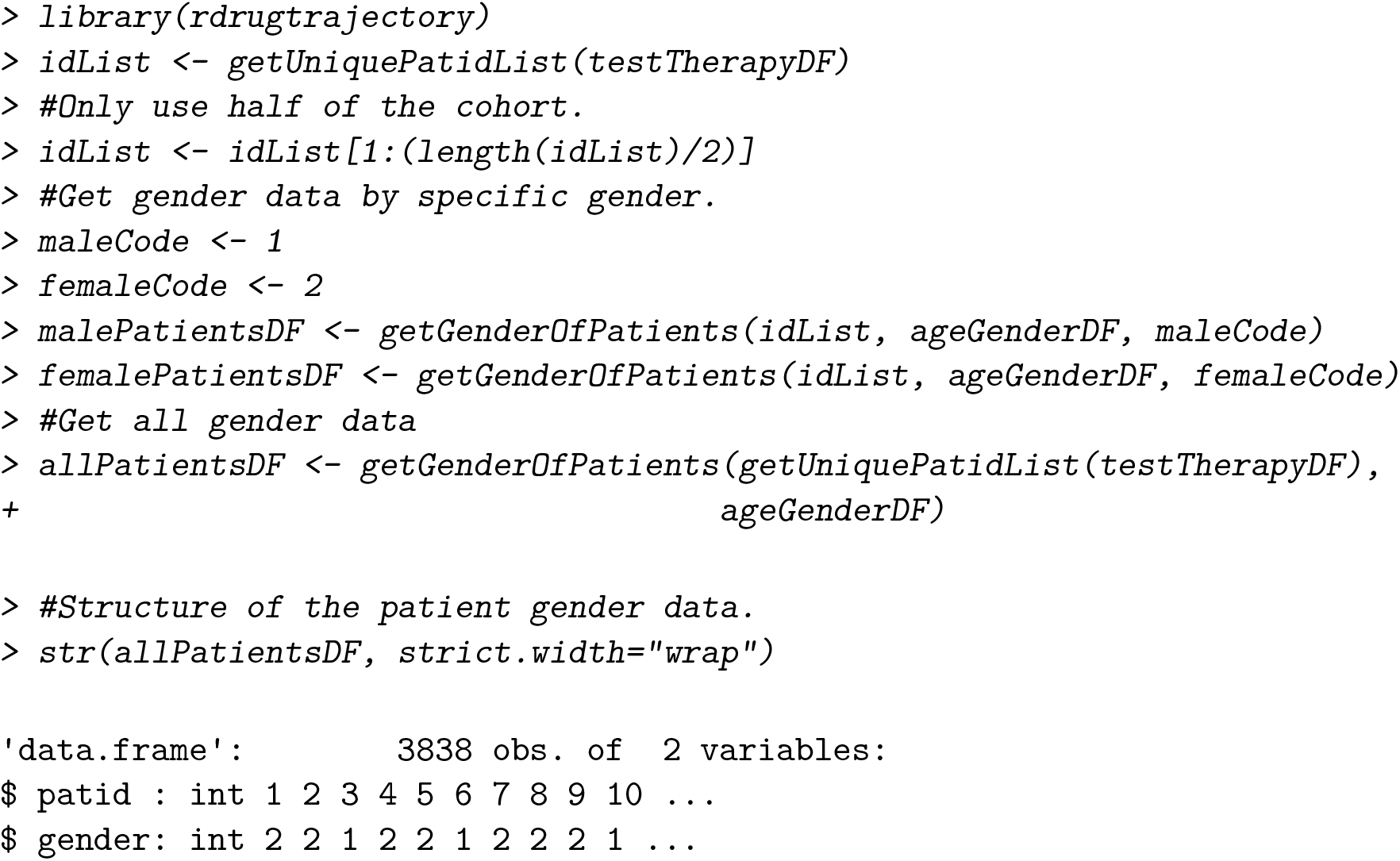

IMD data can be retrieved by combining `getUniquePatidList` and `getIMDOfPatients` functions:

~~~
*> library(rdrugtrajectory)
> idList < - getUniquePatidList(testTherapyDF)
> #Get patients with an IMD score of 1 or 2
> onePatientsDF <- getIMDOfPatients(idList, imdDF, 1)
> twoPatientsDF <- getIMDOfPatients(idList, imdDF, 2)
> #Get all IMD scores for all patients in testTherapyDF
> allPatientsDF <- getIMDOfPatients(getUniquePatidList(testTherapyDF), imdDF)
> #Structure of the patient gender data.
> str(allPatientsDF, strict.width=“wrap”)*
‘data.frame’: 2123 obs. of 2 variables:
$ patid: int 6 11 16 34 42 44 54 60 63 79 ...
$ score: int 1 3 1 4 1 2 1 5 1 2 ...
~~~

The final example of EHR dataframe manipulation presented here demonstrates how to retrieve all prescription records for patients prescribed a specific prescription treatment. For example, such an operation can be used to retrieve all prescription records for any patient prescribed amitriptyline. In addition, it is also possible to return only prescription records matching specific prescription treatments. Importantly, prescription *prodcodes* can be grouped into lists and used to collect those patients with at least one record that matches an element of that list. This approach is useful if the dose is not relevant to the study or the prescription is dispensed under multiple product names.

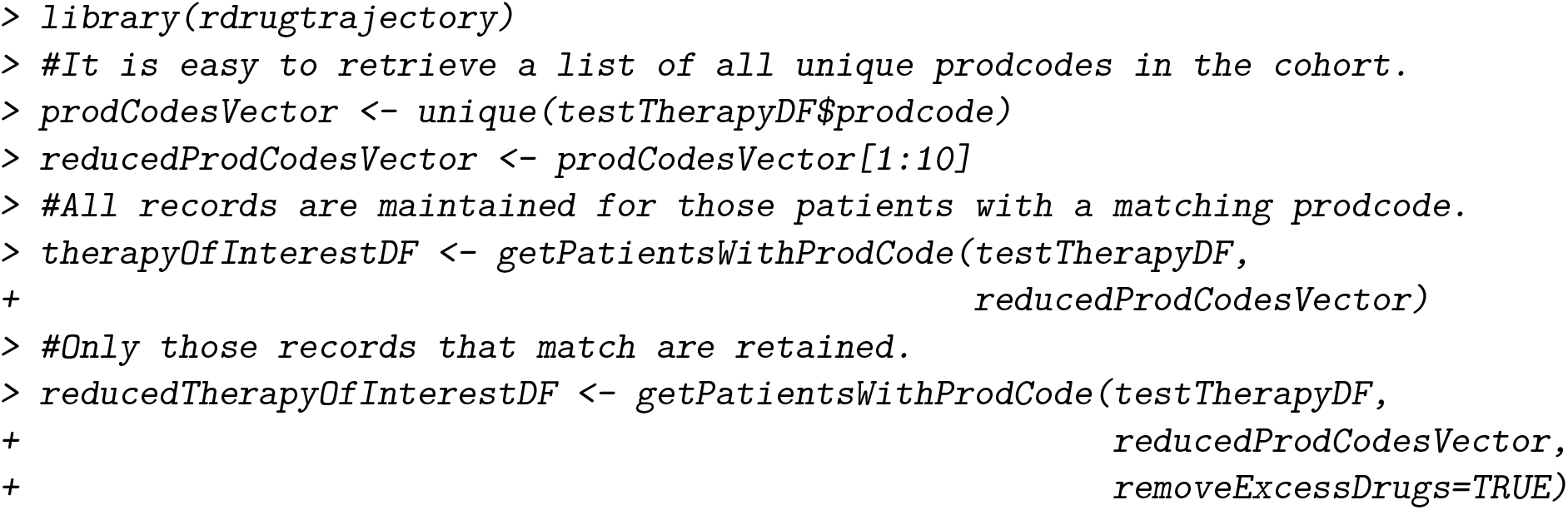

## 3. EHR drug prescription results and discussion

Having briefly demonstrated some basic operation on retrieving patient records by matching EHR dataframes against sets of *patid* values, we move on to showcase several operations available to the user. We begin by presenting examples of cohort prescription summary statistics followed by methods of dataset curating and stratifying by patient groups. We then present examples on how to search for patients prescribed with a first-line treatments, followed by presenting some of these patient groups as sequences of prescriptions. Finally, we demonstrate several examples of building time-lines. For futher examples, please see the Github page and reference manual.

### 3.1. Cohort summmary statistics

#### getEventdateSummaryByPatient

**rdrugtrajectory** can return summary based statistics on patient and cohort level prescription data with `getEventdateSummaryByPatient` and `getPopulationDrugSummary`, respectively. For example, a single patient (via `getUniquePatidList` and [] dataframe subsetting) prescription history returns the patient *patid*, number of prescription events, median number of days between events, fewest number of days between events, the most number of days between events (*maxTime* and *longestDuration* are the same), and record duration (number of days between the first and last prescription event on record):

~~~
*> library(rdrugtrajectory)
> idList <- getUniquePatidList(testTherapyDF)
> resultList <- getEventdateSummaryByPatient(
+ testTherapyDF[testTherapyDF$patid==idList[[1]],])
> str(resultList, strict.width=“wrap”)*
List of 2
$ TimeSeriesList: num [1:6] 336 652 2540 34 42 44
$ SummaryDF :‘data.frame’: 1 obs. of 7 variables:
..$ patid : int 3515
..$ numberOfEvents : int 7
..$ medianTime : num 190
..$ minTime : num 34
..$ maxTime : num 2540
..$ longestDuration: num 2540
..$ recordDuration : int 3648
- attr(*, “class”)= chr “EventdateSummaryObj”
~~~

#### getPopulationDrugSummary

This approach can be extended across the cohort of patients with `getPopulationDrugSummary`. The returning *PopulationEventdateSummary* S3 object is a list of three elements. The first element is the *SummaryDF* dataframe derived from calling `getEventdateSummaryByPatient` per patient, with the set of statistics retrievable through the accompanied *patid*. The second element is the *TimeSeriesList*, which holds a vector per patient of the number of days between consecutive prescription events. Vectors can be accessed using the *patid* element name:

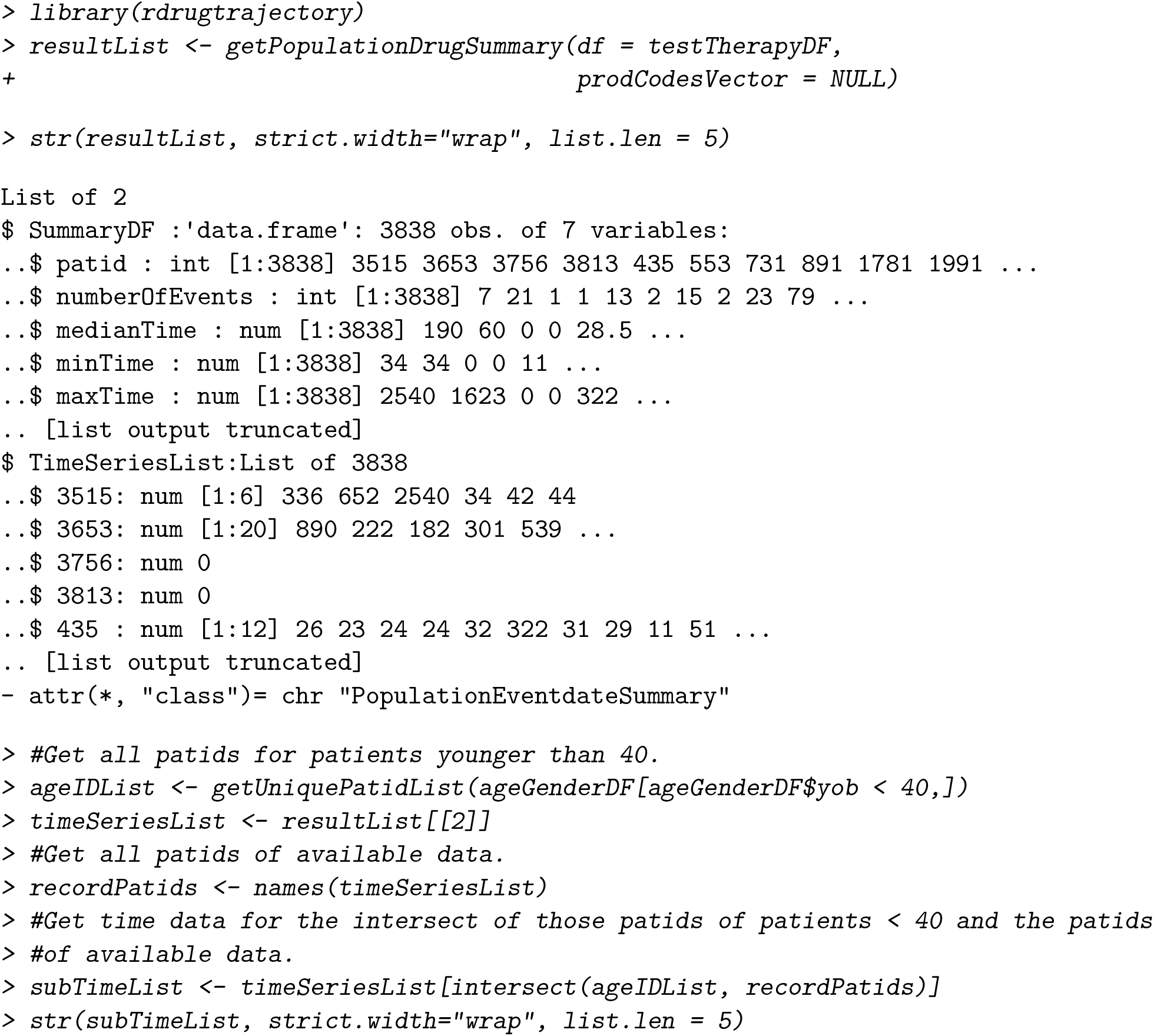
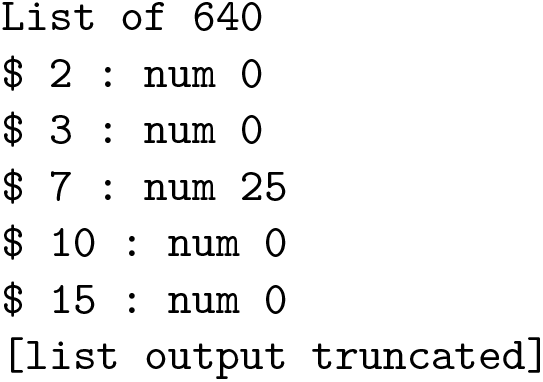

### 3.2. Curating drug prescription records

There is no direct link between a prescription event and a *medcode* in the CPRD data. The relationship between the two can be inferred from the event dates of the prescription and clinical events, in addition, to information provided by the consultation ID and the prescription issue number.

#### matchDrugWithDisease

**rdrugtrajectory** provides several methods for curating prescription datasets with the aim of establishing a relationship between prescription and clinical events. The `matchDrugWithDisease` function returns a subset of all prescription events with an established relationship between therapy and clinical event. To what degree these patients are included in the search is controlled with a function argument. There are three scenarios: all patients with a record of a specific prescription event and specific clinical event, at any point; all patients with a record of a specific prescription event on the same date as a specific clinical event; and, all patients with a record of a specific prescription event on the same date as a specific clinical event and clear from additional clinical events on that day. One would expect fewer patients as the stringency of the search criteria is increased:

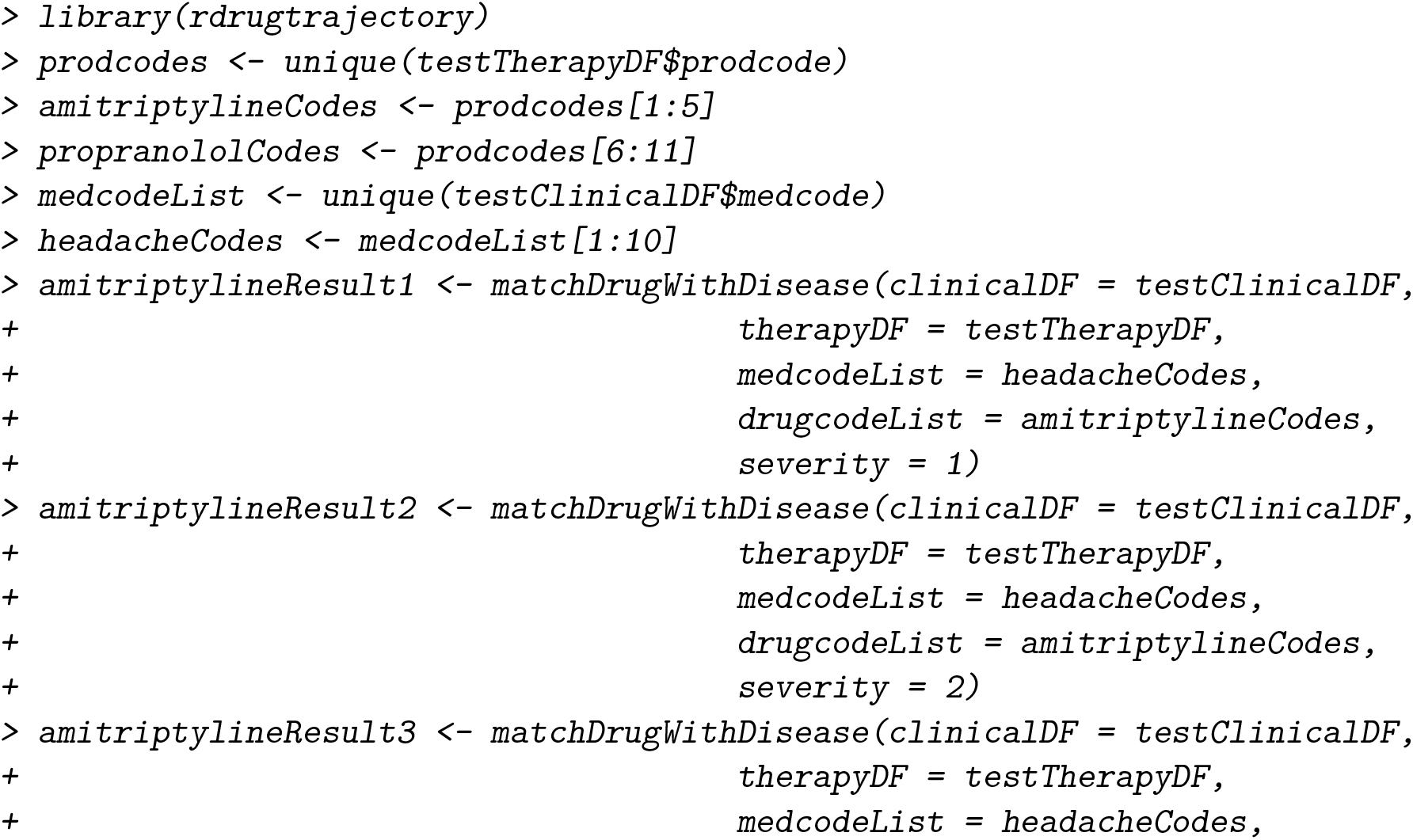
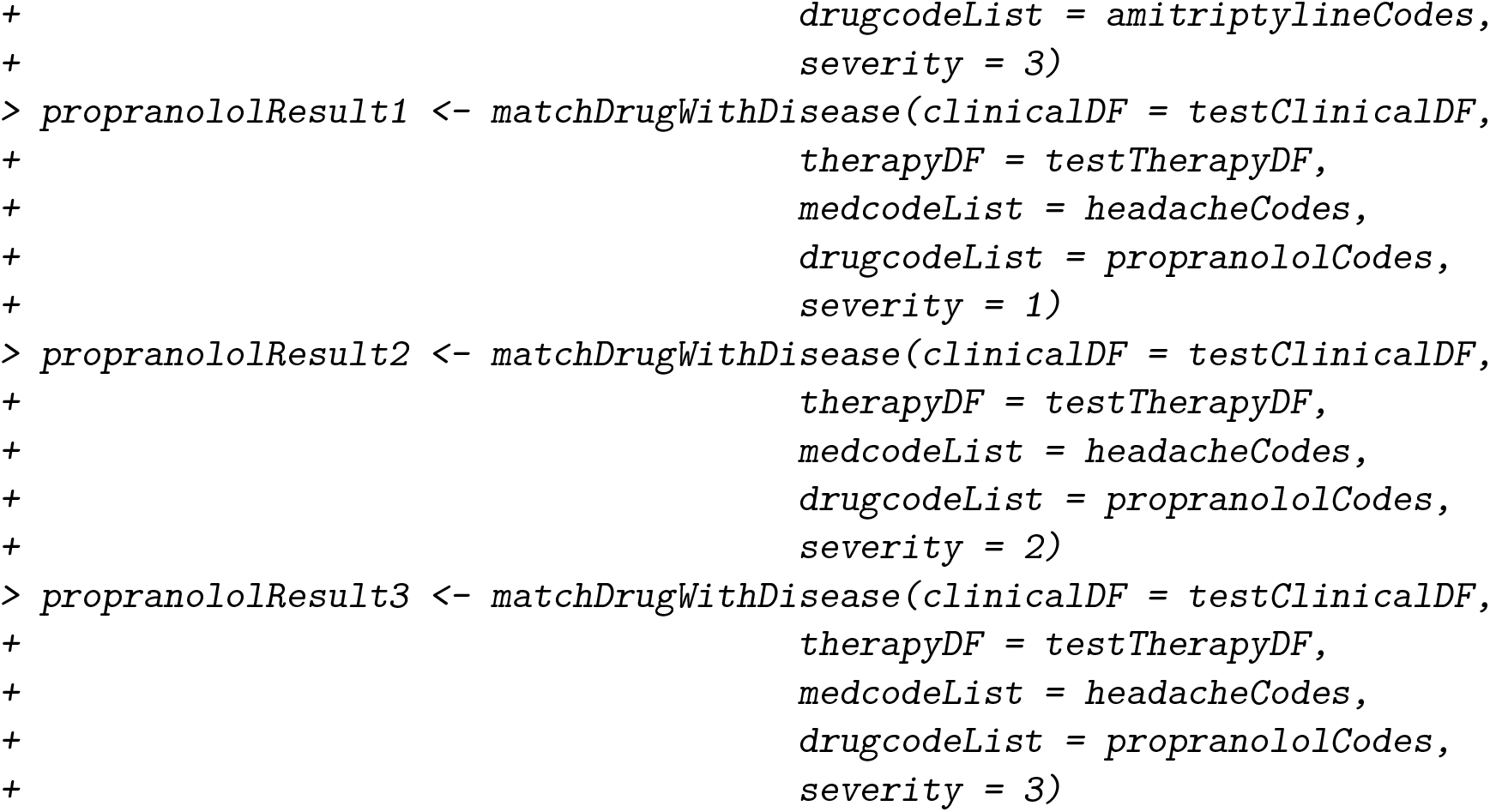

#### getGenderOfPatients

The example presented, demonstrates how to identify patients prescribed amitriptyline and patients prescribed propranolol (there is patient overlap, easily controlled for by subsetting) whilst controlling for clinical overlap with or without consideration for off topic clinical events. With the identified patients, we can, for example, stratify by gender:

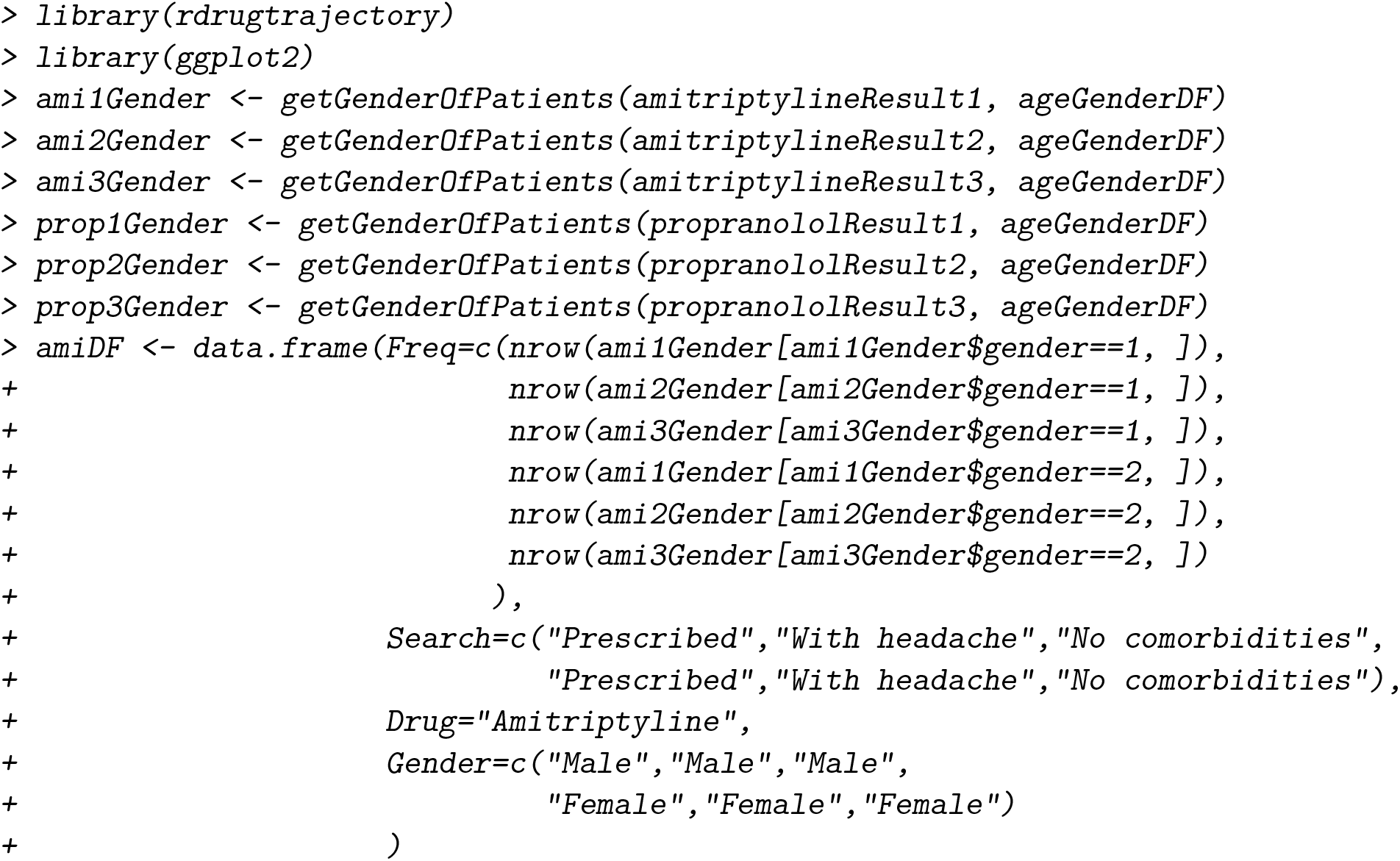
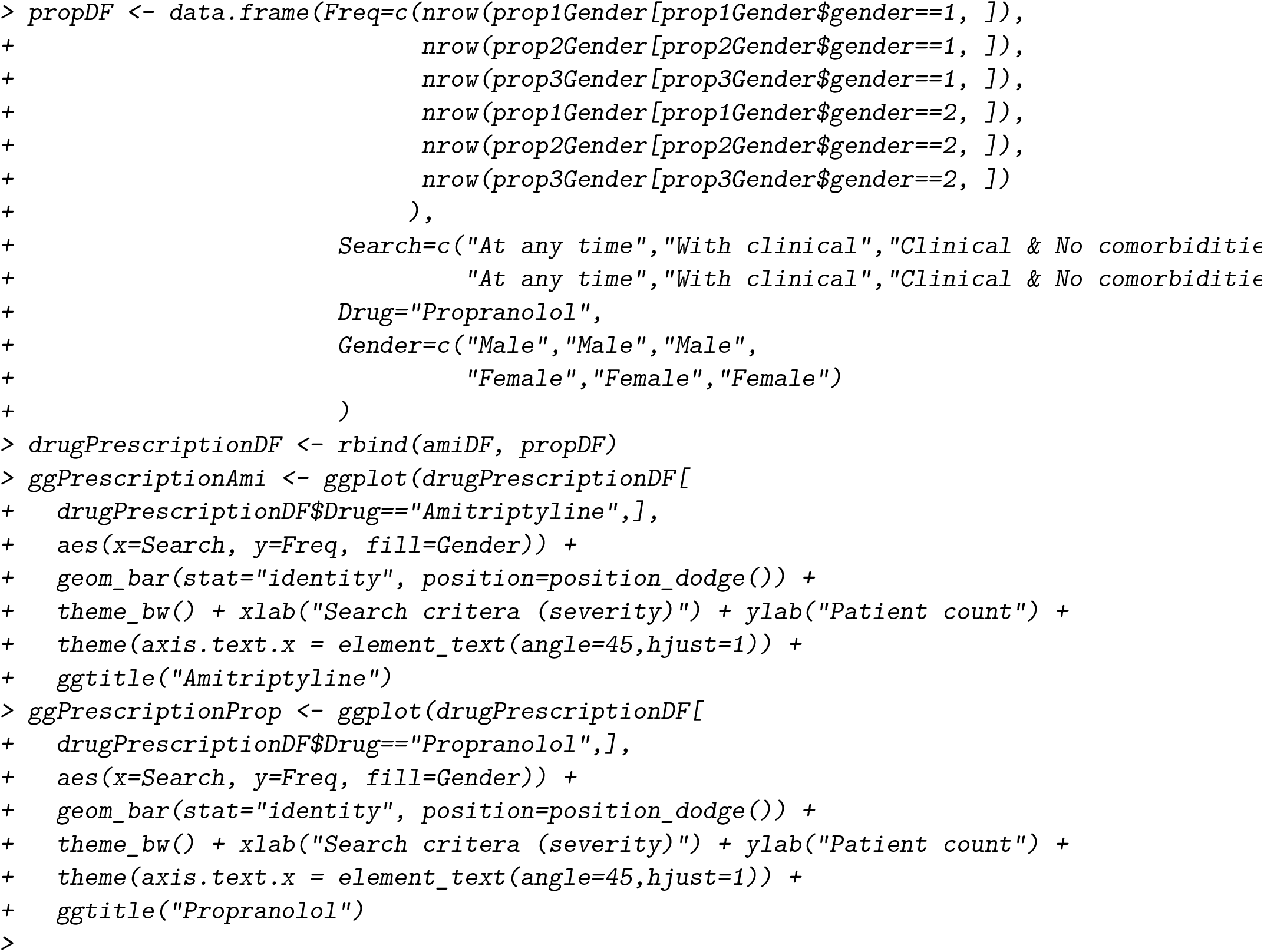

Filtering through prescription events can also be controlled by a date range. For example, if one was calculating the number of patients prescribed amitriptyline per year from 2000 to 2004 and matched to a headache event, one can apply a date range:

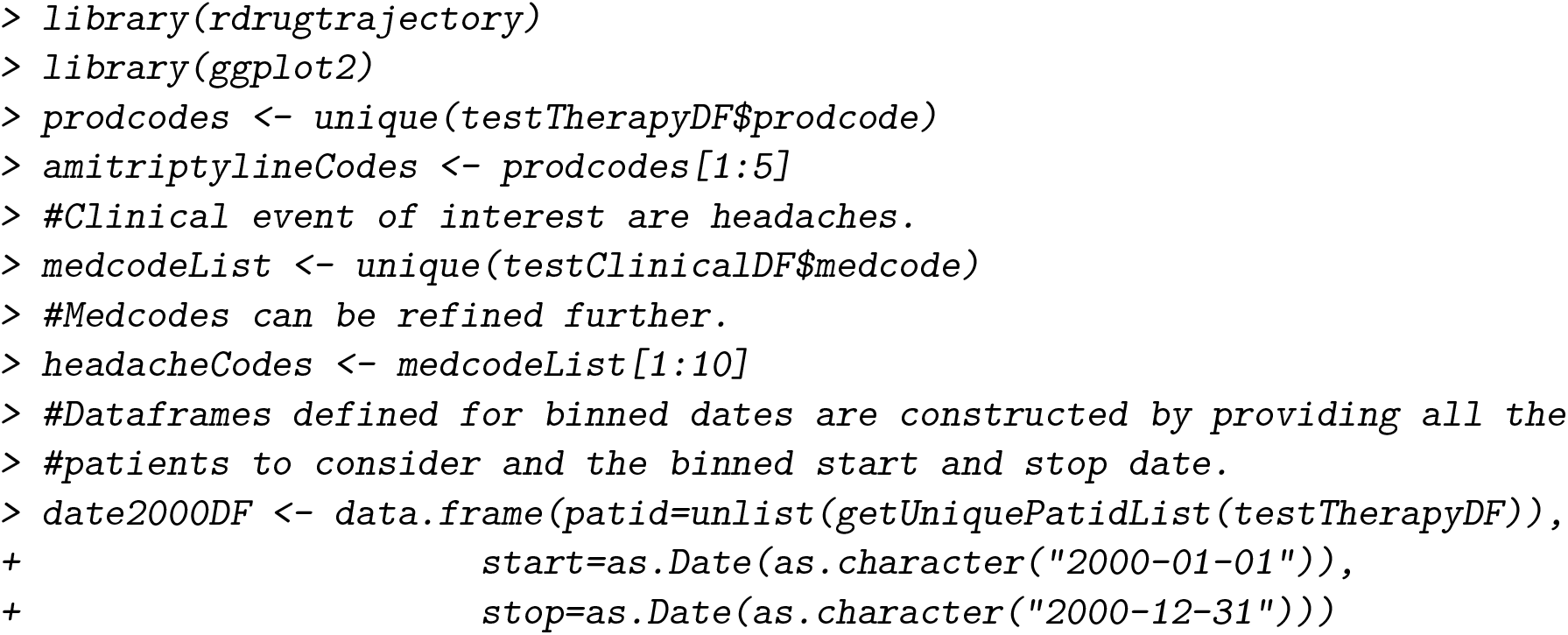
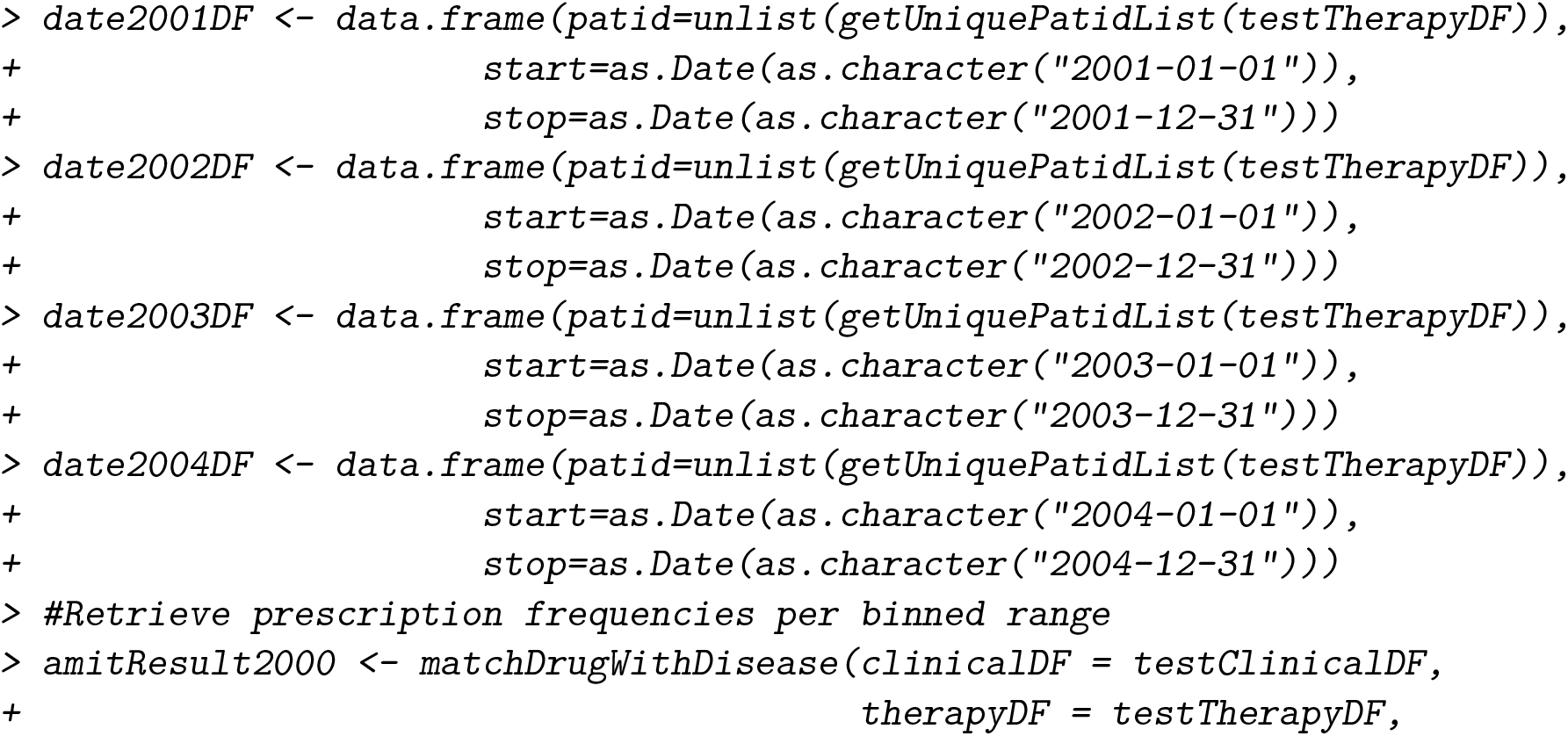
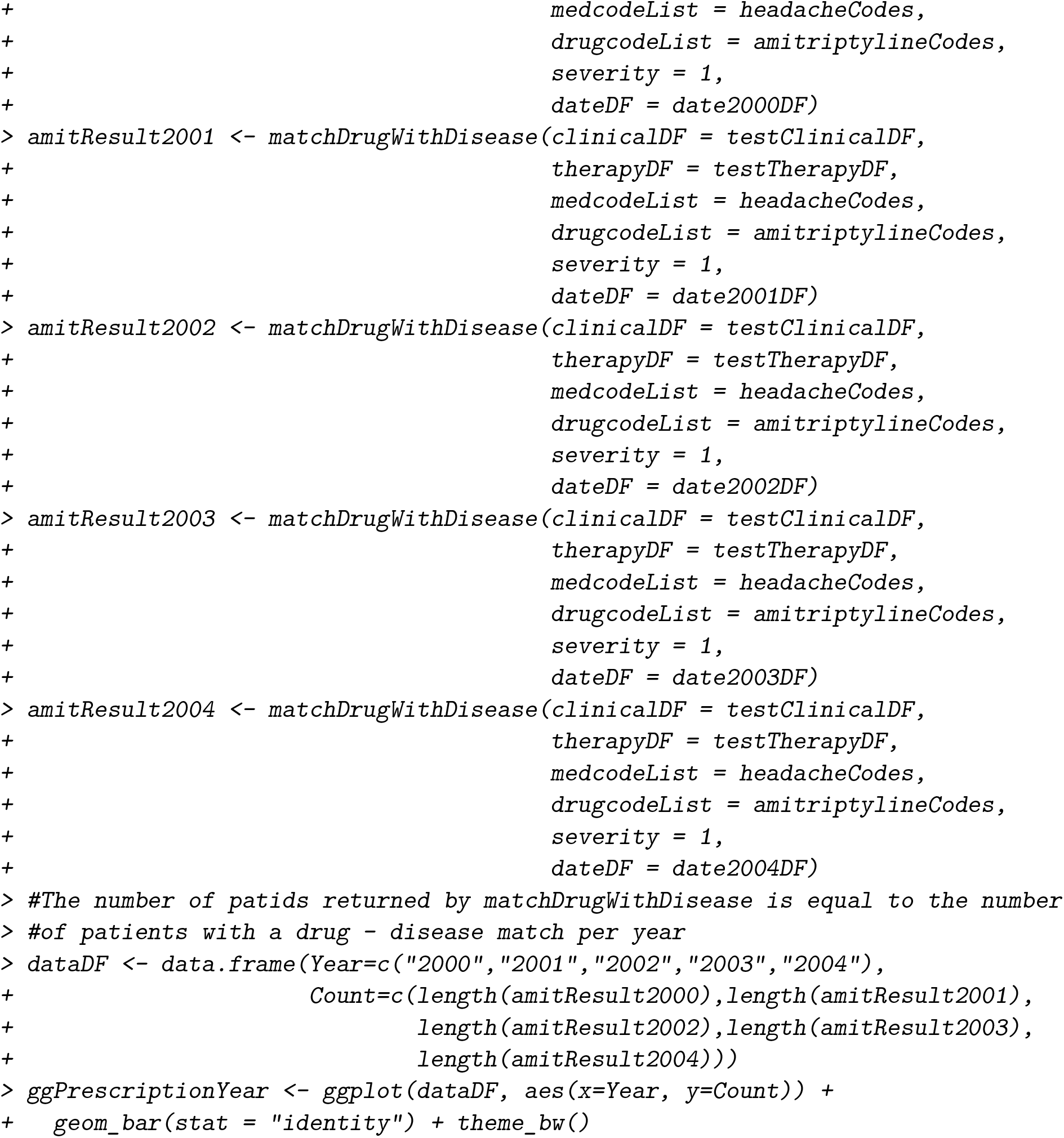

#### getPatientsWithFirstDrugWithDisease

Unlike `matchDrugWithDisease` which retrieves patients with a prescription event matching clinical criteria at any time within a CPRD EHR record, `getPatientsWithFirstDrugWithDisease` identifies patients with a first prescription event that matches a desired clinical event. Please note, care must be taken when searching for medication with off-label uses. For example, beta-blockers are frequently prescribed to treat hypertension and arrhythmia, however, the beta-blocker propranolol is also prescribed to treat migraine. Without in depth analysis into the patient history, patients propranolol with records for hypertension or arrhythmia in addition to migraine on a matching *eventdate* with the first propranolol prescription, could result in a misleading disease-drug association. In cases where a health care professional suggests a change in the patient’s lifestyle choices, that patient may have several clinical events free from prescriptions before the first prescription of interest is prescribed. Using basic subsetting one can calculate the number of clinical events before the patient’s first prescription intervention (Figure 3 A). Further more, we can stratify patients into subgroups (Figure 3 B):

**Figure 3:**
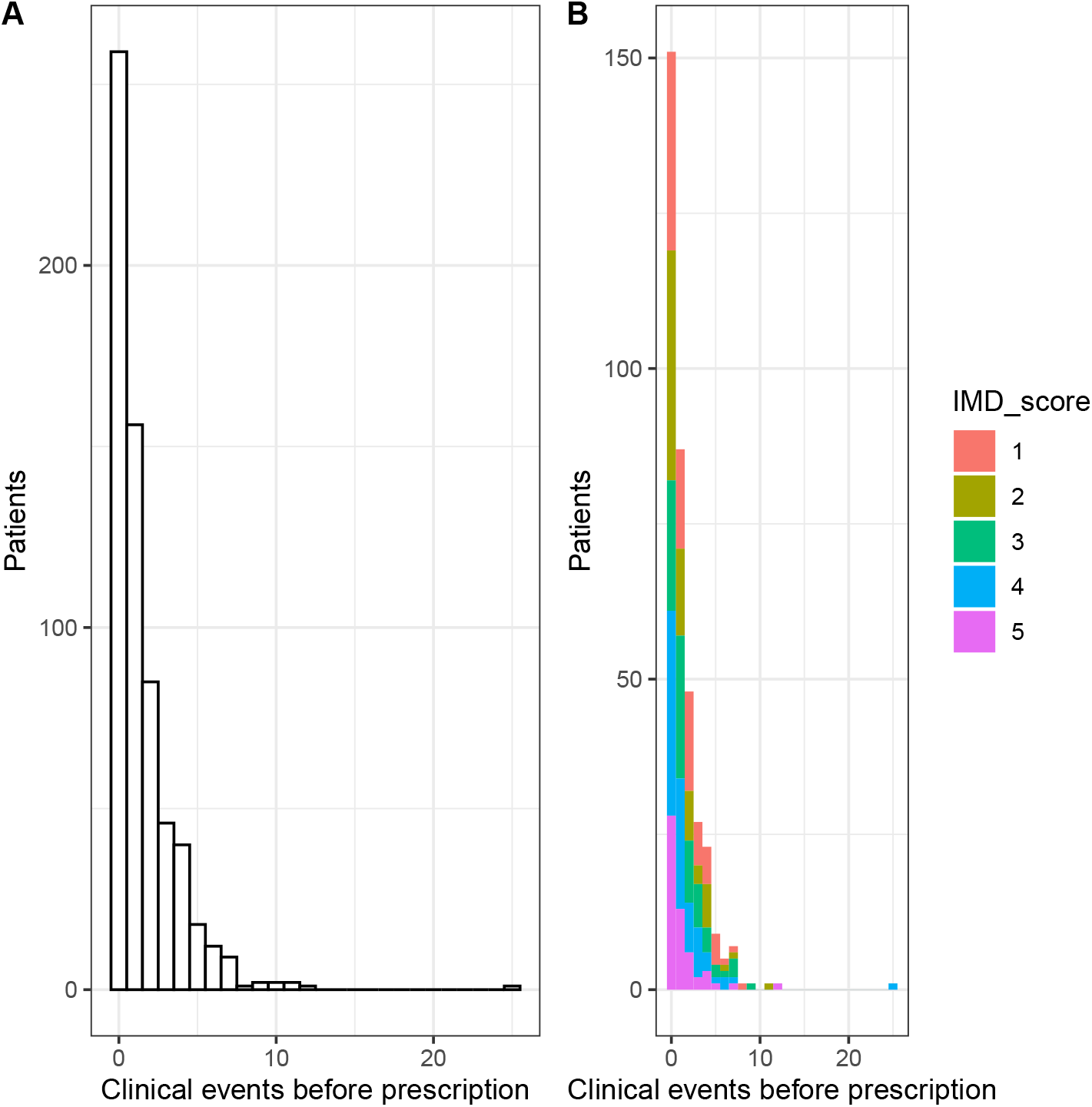
The number of clinical events before the first treatment across the whole cohort (A), and by IMD score (B).

**Figure 4:**
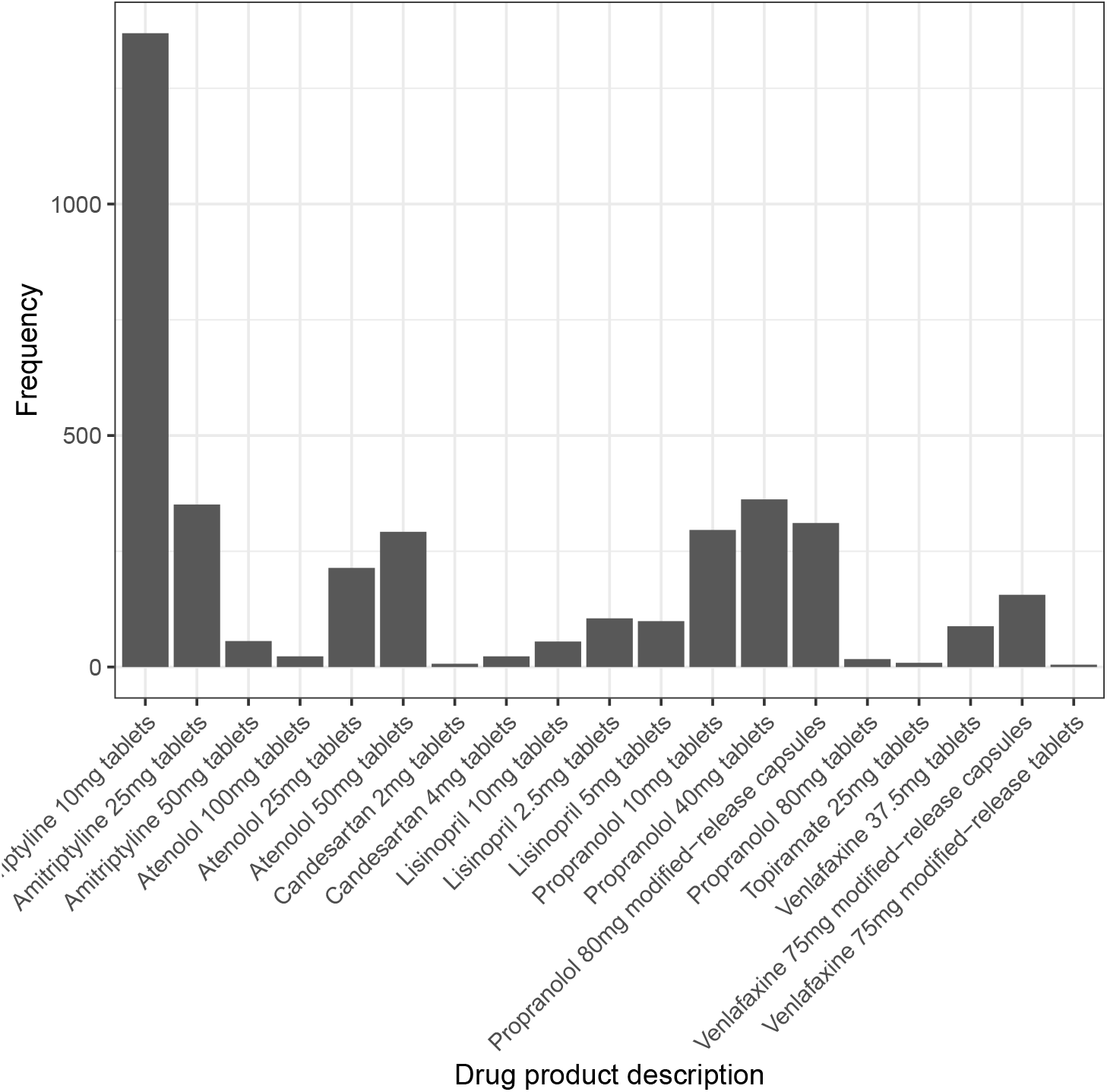
The frequency of first line treatment prescription.

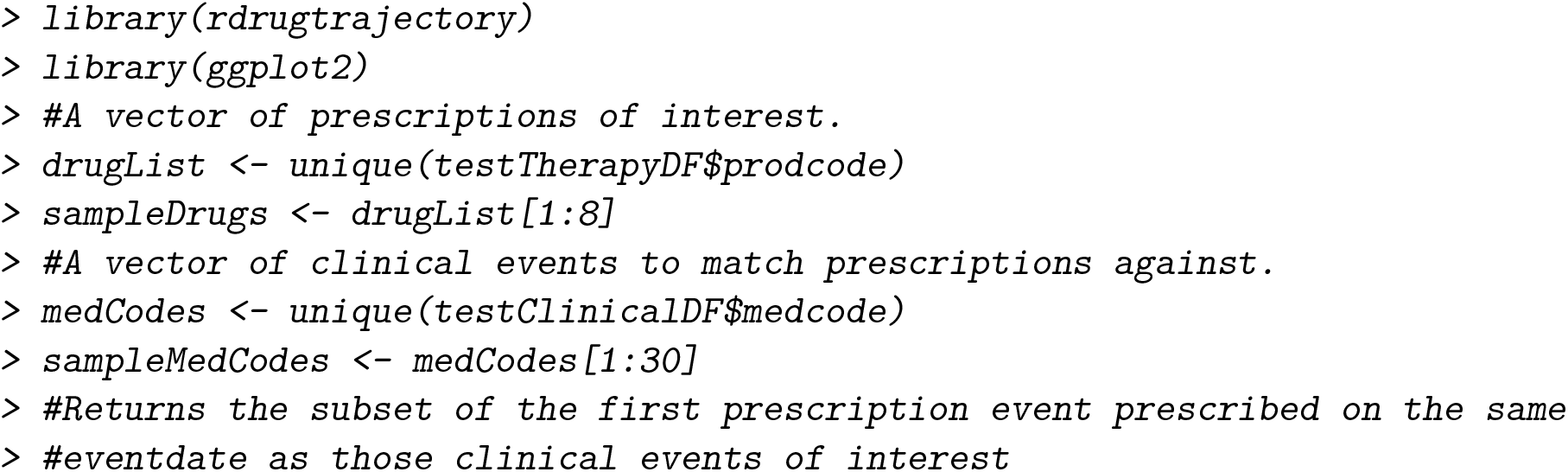
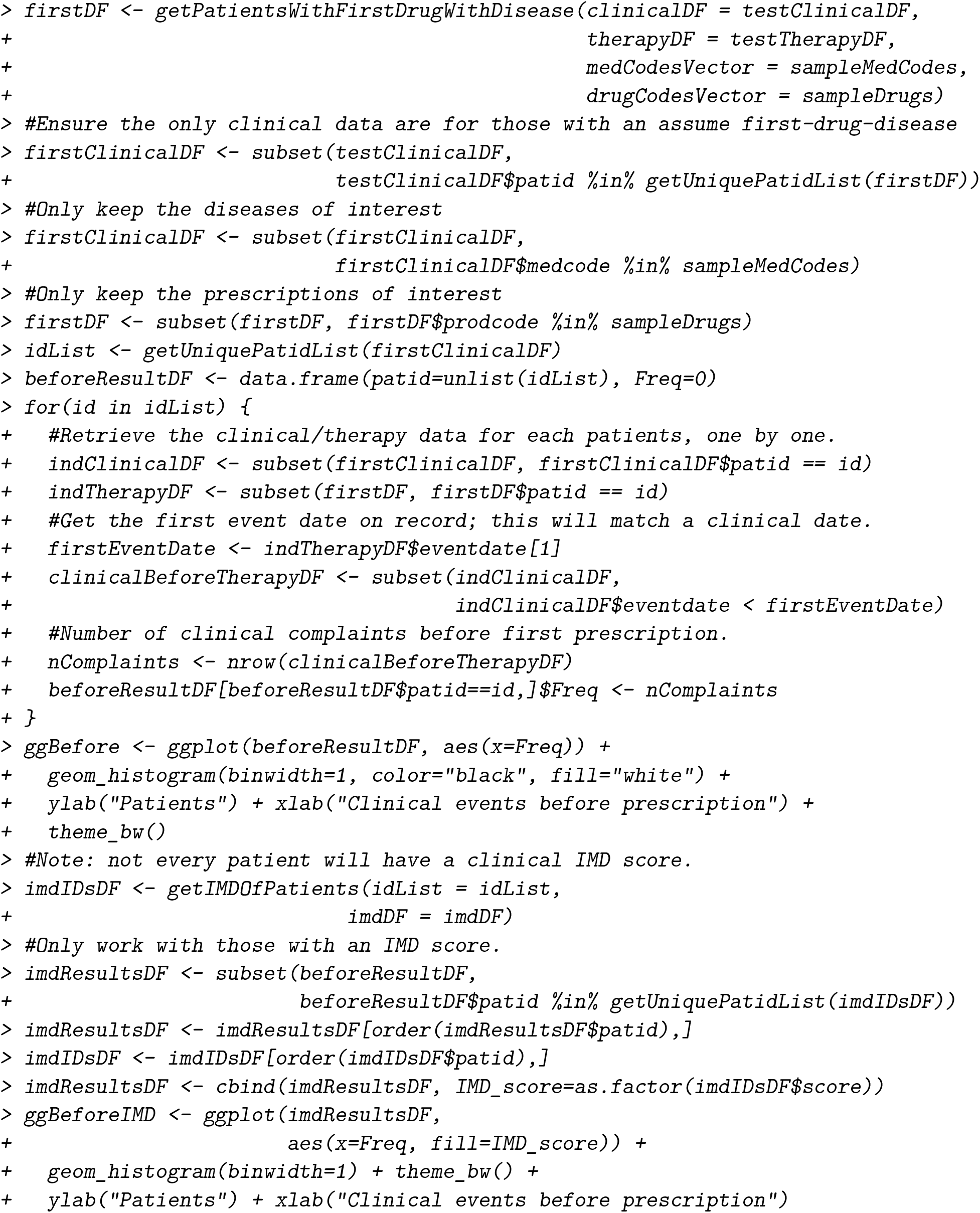

#### getMultiPrescriptionSameDayPatients

The function `getMultiPrescriptionSameDayPatients` returns all prescription events for those patients prescribed more than two prescriptions on the same date. All events of those patients without a prescription *prodcode* event can be removed. Combining `getMultiplePrescriptionSameDayP` with `getPatientsWithFirstDrugWithDisease` or `matchDrugWithDisease` is useful for filtering patients for specific prescription patterns. For example, to retrieve all patient prescription records if specific prescriptions are (a) never recorded together on the same date and (b) are used as a first line treatment for a given complaint:

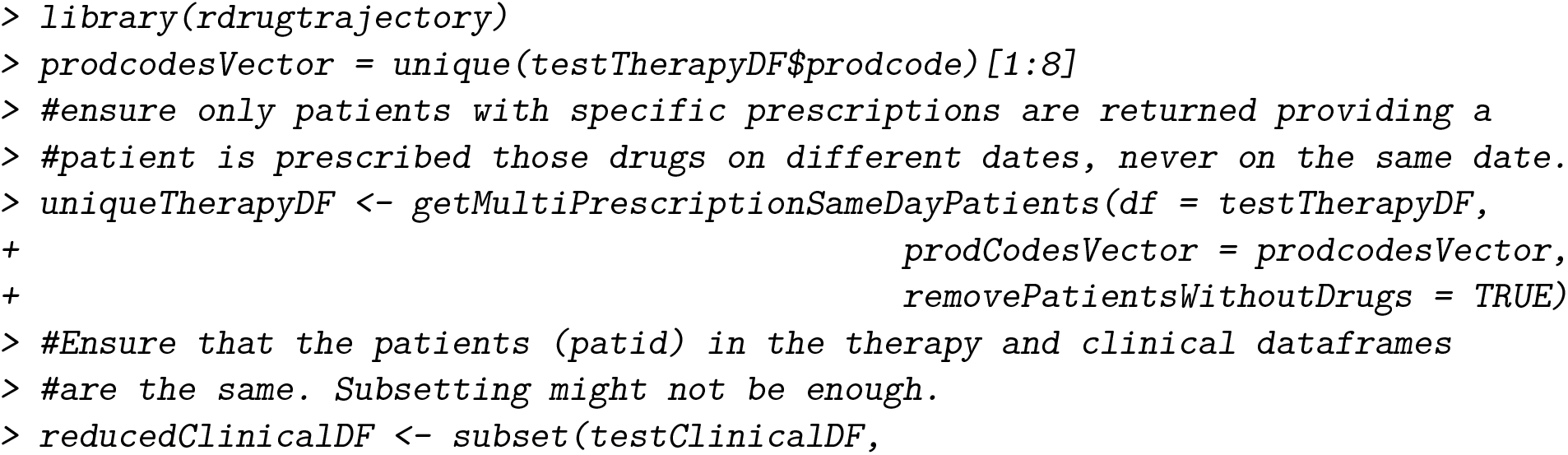
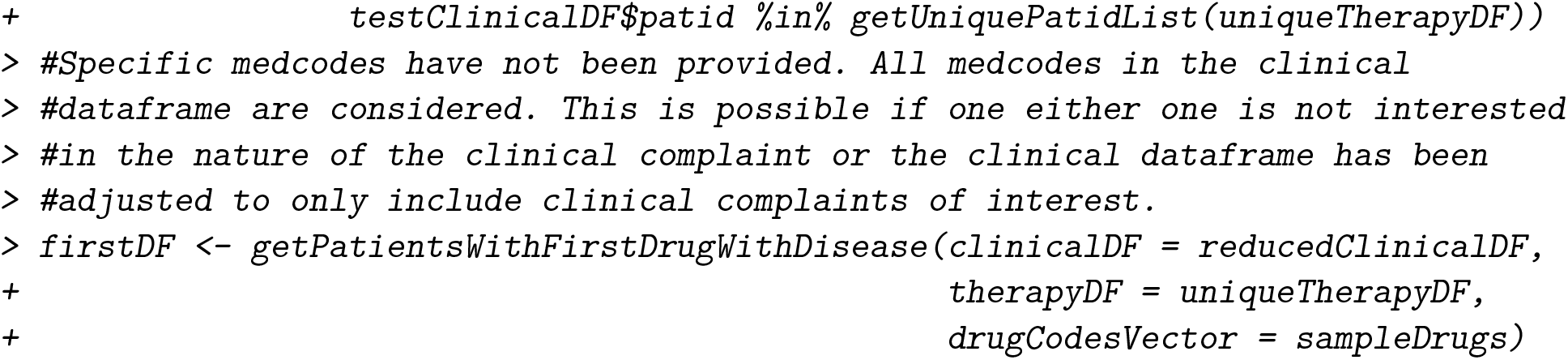

In the above example, patients with more than one prescription on the same date or without a prescription at all (from the set of desired prescription *prodcodes*) were removed from the cohort. This reduced the number of patients from 3838 patients to 2930. Next, only those patients with a first line treatment (first prescription event on the same date as a clinical event) were kept, reducing the sample size to 587 patients.

#### removePatientsByDuration

Longitudinal EHR cohort studies often requires careful time-related consideration. Currently, **rdrugtrajectory** presents two functions that identify prescription records of patients that match two time constraints. The first, `removePatientsByDuration`, removes all patients with prescription events that are no more than *n* years between consecutive events or removes patients if the duration between the first and last prescription event on record is less than *n* years.

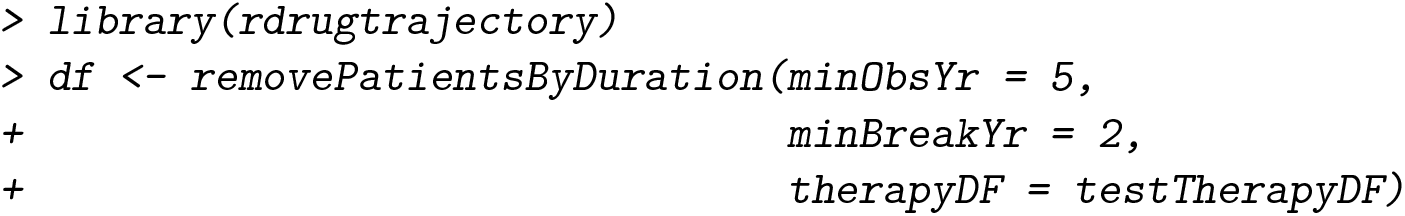

##### getBurnInPatients

The second time-related function, `getBurnInPatients` identifies all patient prescription records with at least *n* days free from prescription events before a specific prescription event. This is useful if one requires a period of time free from prescription intervention before a given prescription event:

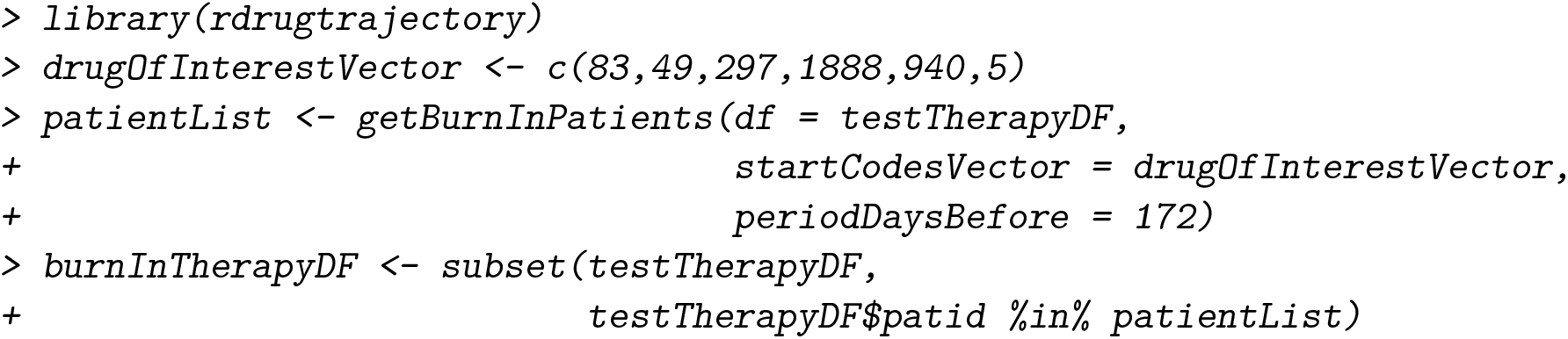

In the above example, from a cohort of 3838 patients, 426 patients had a period of up to 172 days free from of prescription events before the first prescription *prodcode* specified via the *startCodesVector* argument. The functionality relies on the patient having prescription events before the burn-in period (required to define whether the patient had a CPRD record early enough before the burn-in period began). For example, this patient had over three years of prescription events before the prescription of interest (from *2003-05-29* to *2007-10-17* with over 172 days free from exposure before the prescription event of interest *prodcode* 297:

~~~
*> head(burnInTherapyDF[burnInTherapyDF$patid == 332412,], n=9)*
[1] patid eventdate prodcode consid issueseq
< 0 rows> (or 0-length row.names)
~~~

### 3.3. First drug prescriptions

#### getFirstDrugPrescription

A patient’s first prescription event on CPRD record can be identified by supplying `getFirstDrugPrescriptio` with a list of prescription *prodcodes*. The functions returns `FirstDrugObject`, an R S3 object of type List. Only the first prescription event to match anyone one of the prescription *prodcodes* provided is identified. The first element of `FirstDrugObject` contains a named list of *patid* vectors. Each vector contains the *patids* of all those patients that share the same first prescription *prodcode*. The list element is named after the corresponding prescription *prodcode*. The second element in `FirstDrugOject`, like the first, is a list of `Date` vectors, each named after the corresponding prescription *prodcode*. Each `Date` vector contains the *eventdate* of the prescription event for the patient identified by the *patid* in the identical position of the preceding List. The third list element contains a table of prescription frequencies for each first prescription *prodcode* on record. The *prodcode* is accompanied by a product description providing a file of CPRD prescription products has been provided. Below we demonstrate how to retrieve information on first-line treatment:

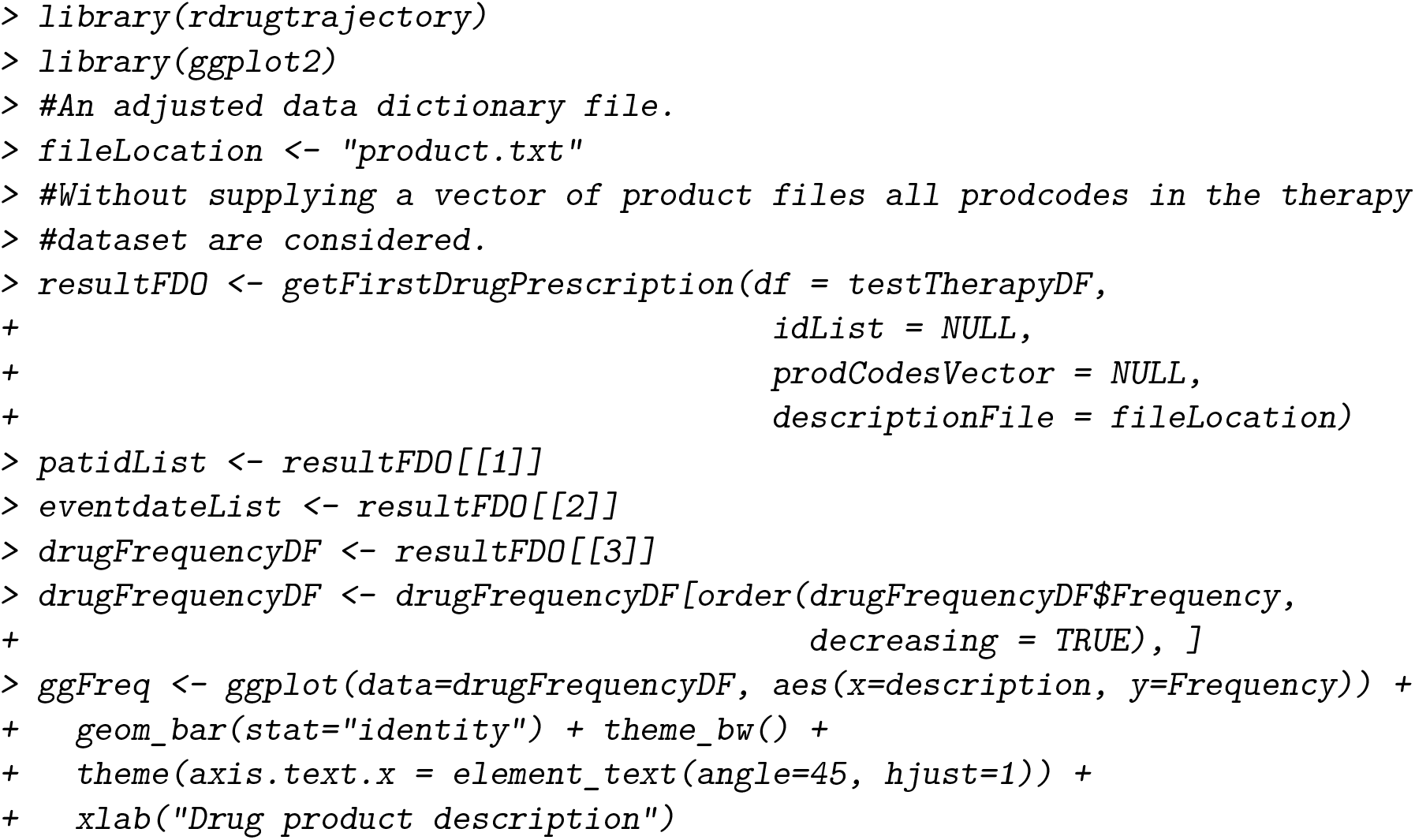
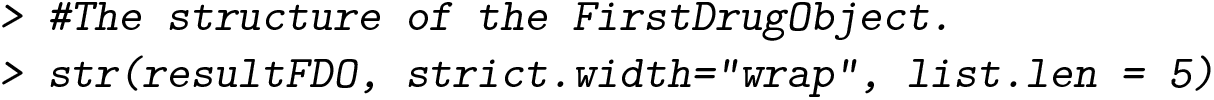

#### getAgeGroupByEvents

In the next example we explore stratifying first-line prescription events by patient characteristics, such as, age, gender, IMD, and number of *medcodes* (for instance, by comorbidities) or *prodcodes* (for instance, to separate those patients by additional prescriptions), or by any additional clinical event retrieved using *CPRDLookups*.*R* **?. rdrugtrajectory** provides several utility functions to stratify patients (see reference manual for further information). The function `getAgeGroupByEvents` calculates the number of first-line prescription events by patient age. By specifying a set of *patids* and *eventdates* from the `FirstDrugObject`, we can calculate the number of first-line prescriptions by age-group for patients linked with a specified medical condition:

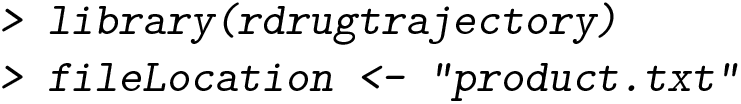
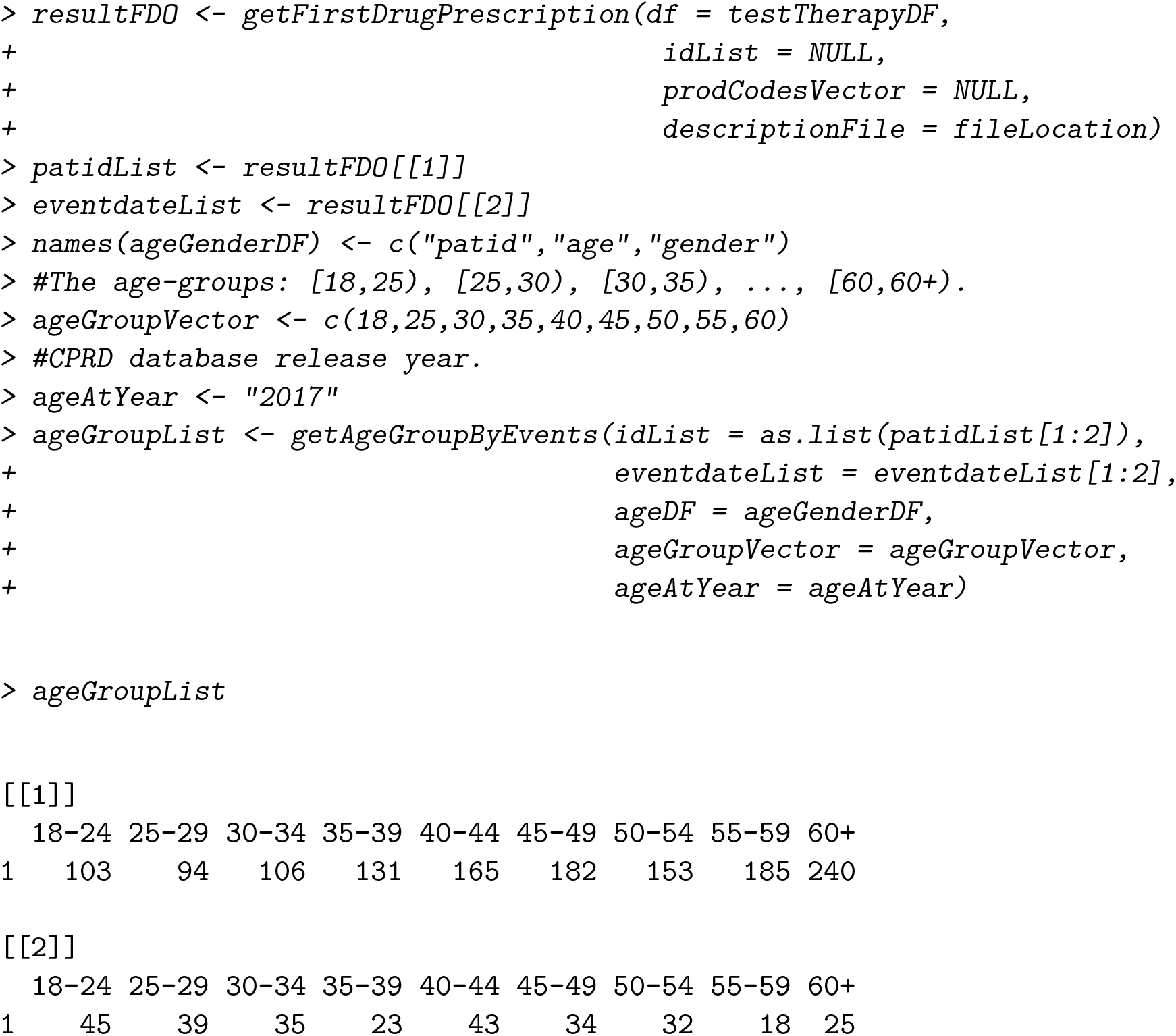

In the above example, the age of each patient (*ageDF*) was provided using year-of-birth calculated against the release year of the CPRD Gold database (explained above). By providing the database release year (in *ageAtYear*) and the first prescription *eventdate* (in *eventdateList*), the age of each patient is adjusted against the prescription *eventdate* year. Finally, by using a list slice on *idList* and *eventdateList*, (individual prescriptions can be specified using their *prodcode*, for example, `eventdateList$’105’`), first prescription prescriptions frequencies by age-group are retrievable (Figure 5).

**Figure 5:**
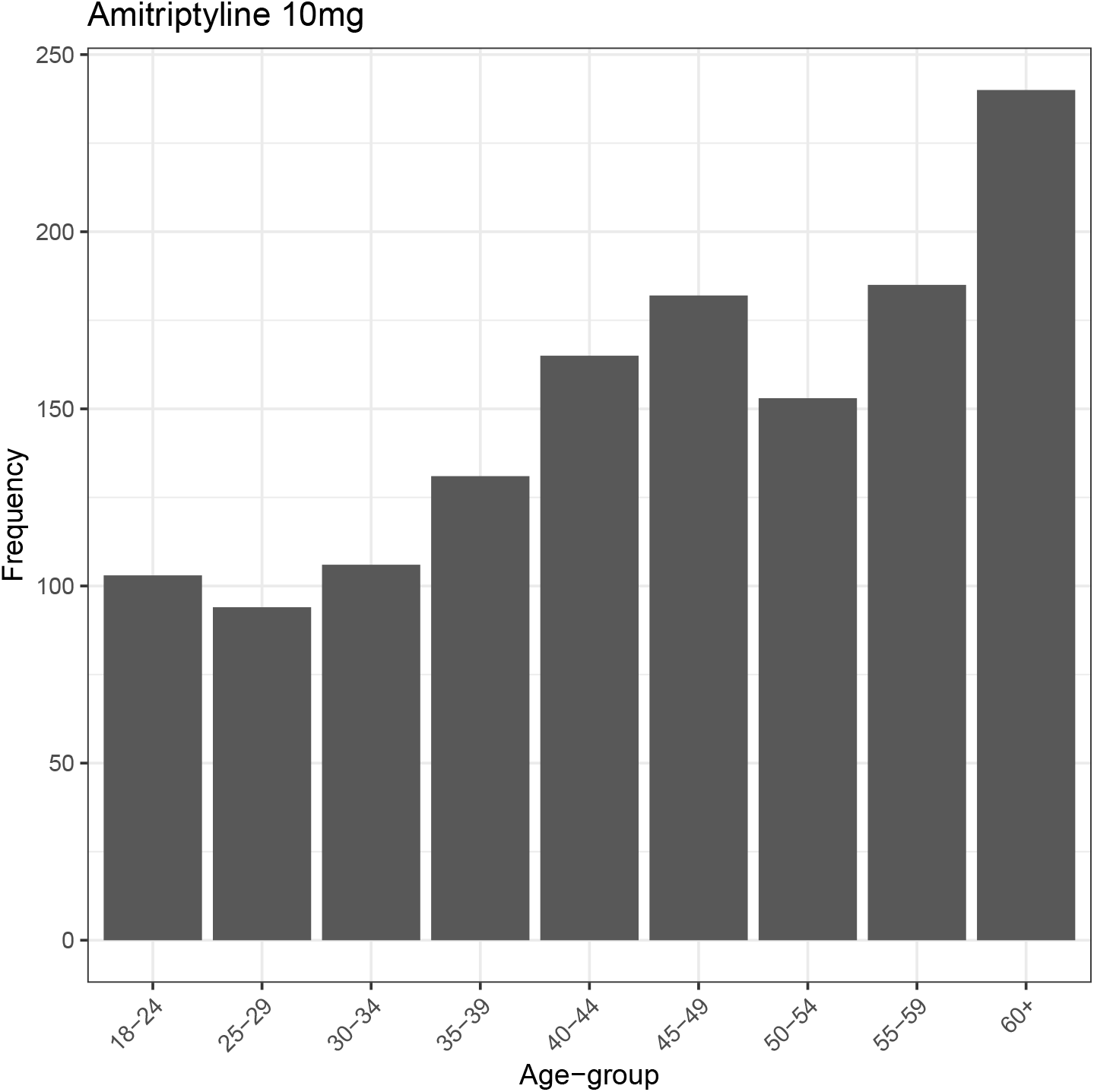
The distribution of Amitriptyline 10mg as a first-line treatment by age-group.

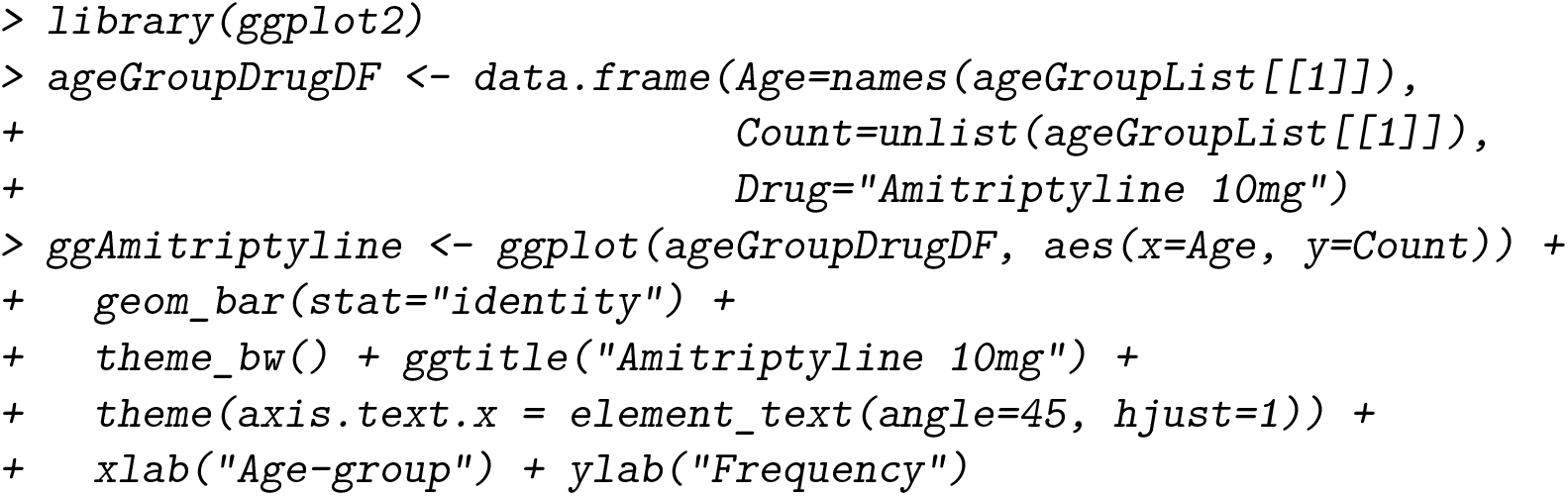

### 3.4. Prescription sequences

#### mapDrugTrajectory

Identifying patient prescription trajectories in longitudinal EHRs remains our biggest motivator behind the development of **rdrugtrajectory**. Therefore, we developed `mapDrugTrajectory` to identify the chronological of patient prescription events. We restrict the calculation to only look for prescription *prodcodes* as supplied to `groupingList` as a named list (named *prodcode* vectors). The required number of grouped-prescription events is defined by specifying the `minDepth` and the number of those changes to display is controlled by `maxDepth` maximum number. By keeping `minDepth` and `maxDepth` the same, only patients with a valid number of prescription changes are displayed (Figure 6 (A) and (C)). Patient records with fewer than `minDepth` number of changes to prescription sequences are ignored (Figure 6 (B)). For further information please refer to the reference manual.

**Figure 6:**
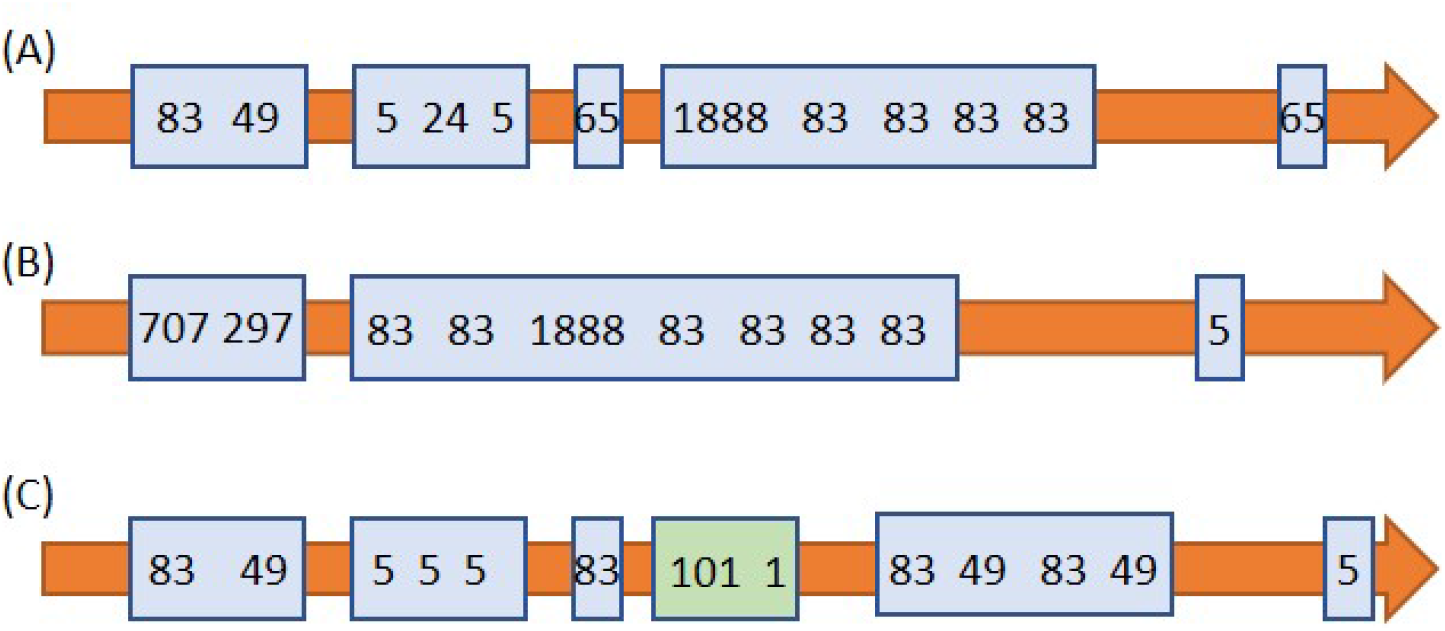
The distribution of grouped *prodcodes* across three patients. (A) Five groups of valid prescription *prodcodes*, (B) only three groups, (C) five valid groups, in addition to *prodcodes 101* and *1* which are ignored.

**Figure 7:**
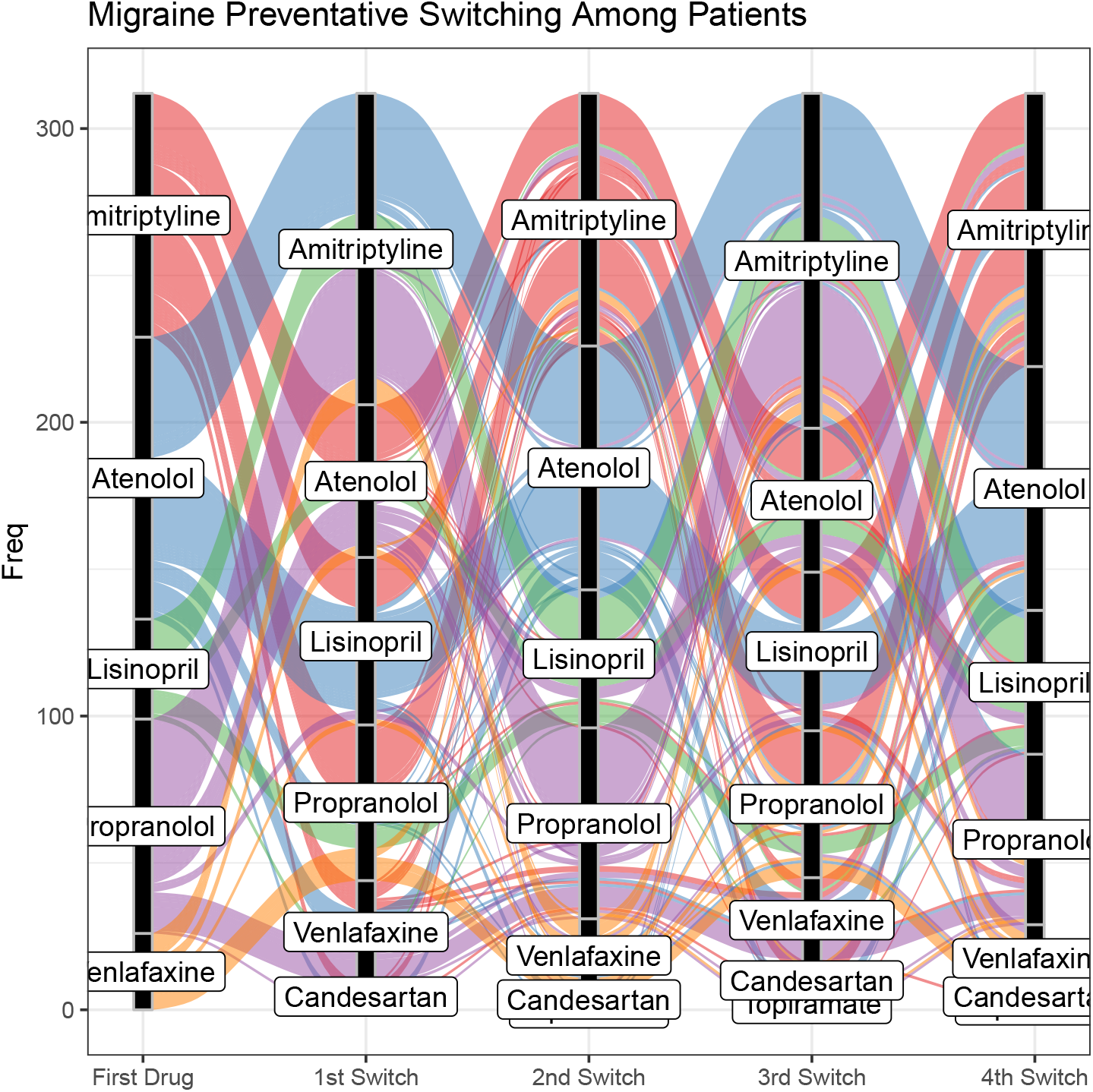
Prescription pattern switching of seven different migraine preventatives. A patient required a a minimum of five changes in prescriptions (including the initial prescription) and, equally, the display was set to five changes in prescription.

**Figure 8:**
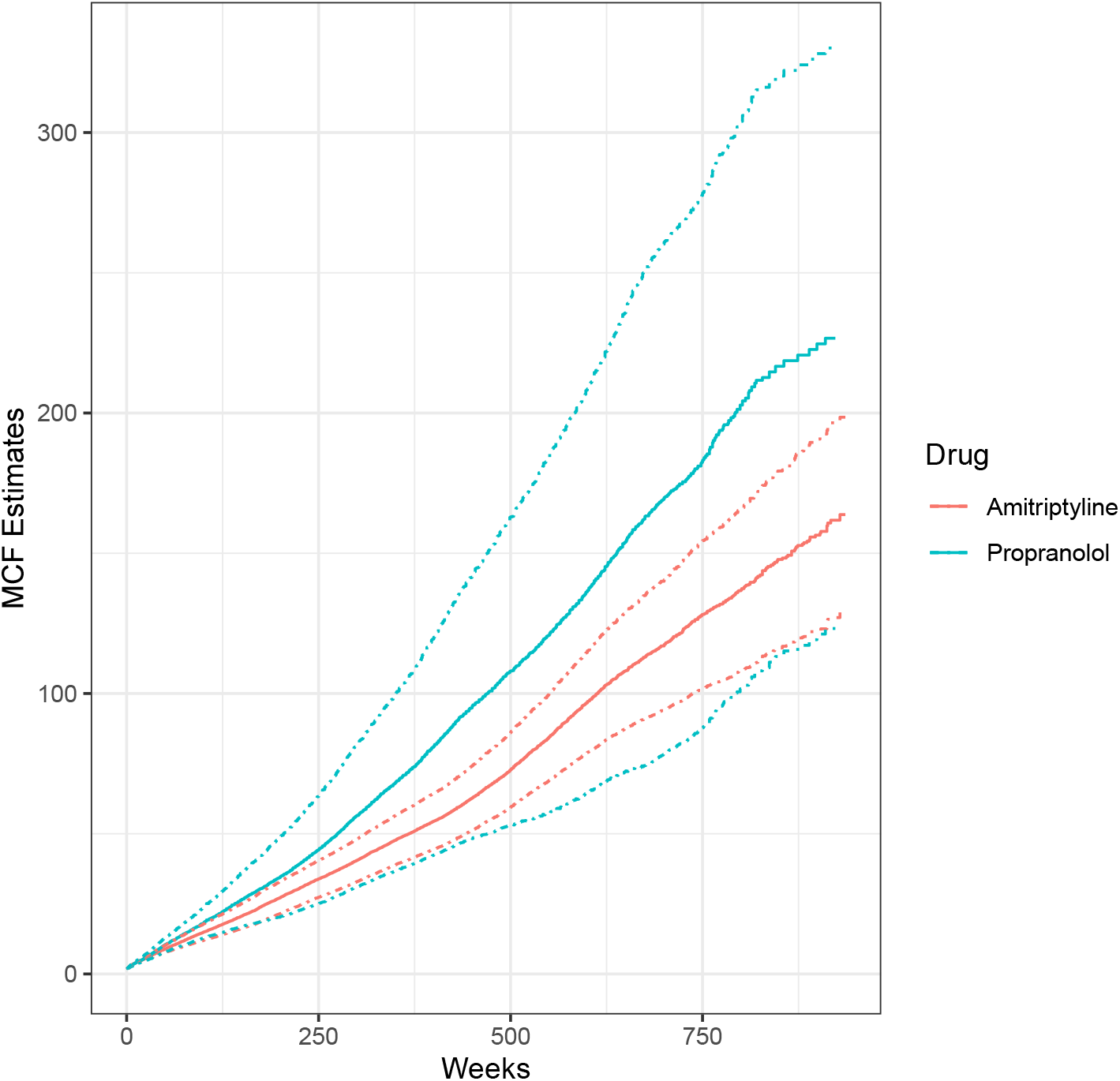
MCF of drug prescriptions of patients with a first drug prescription for either amitriptyline or propranolol, stratified by gender. The dotted lines indicate a 95% confidence interval.

**Figure 9:**
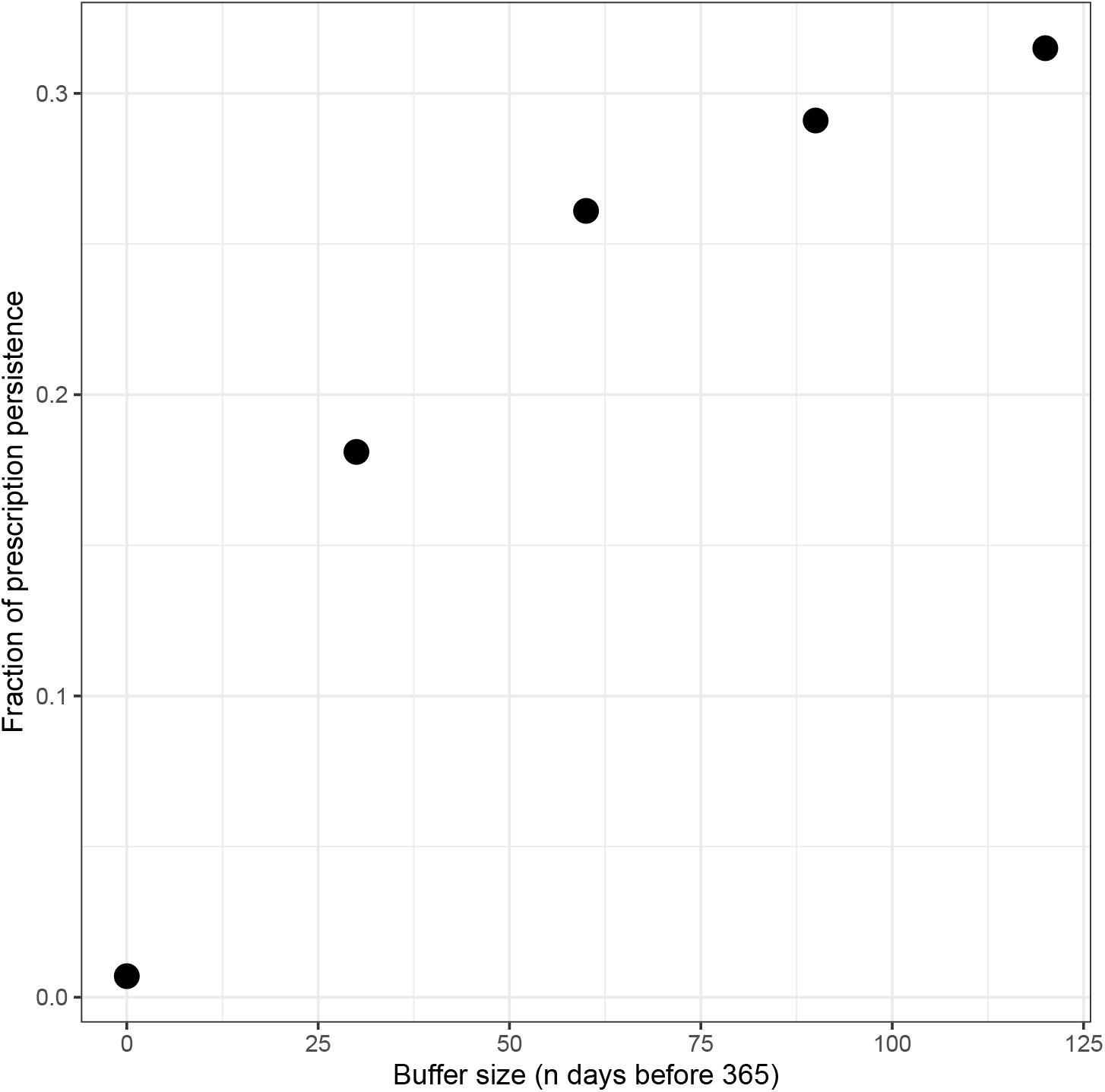
The fraction of prescription persistence adjusted by a `buffer` number of days before a calender year. As the `buffer` approaches the value of `duration` the fraction approaches 1.

In the code below, `mapDrugTrajectory` returns patients with at least first five grouped prescriptions. *prodcodes* that have not been grouped are ignored. Duplication of *prodcodes* (those from the same group) do not count as a change in treatment:

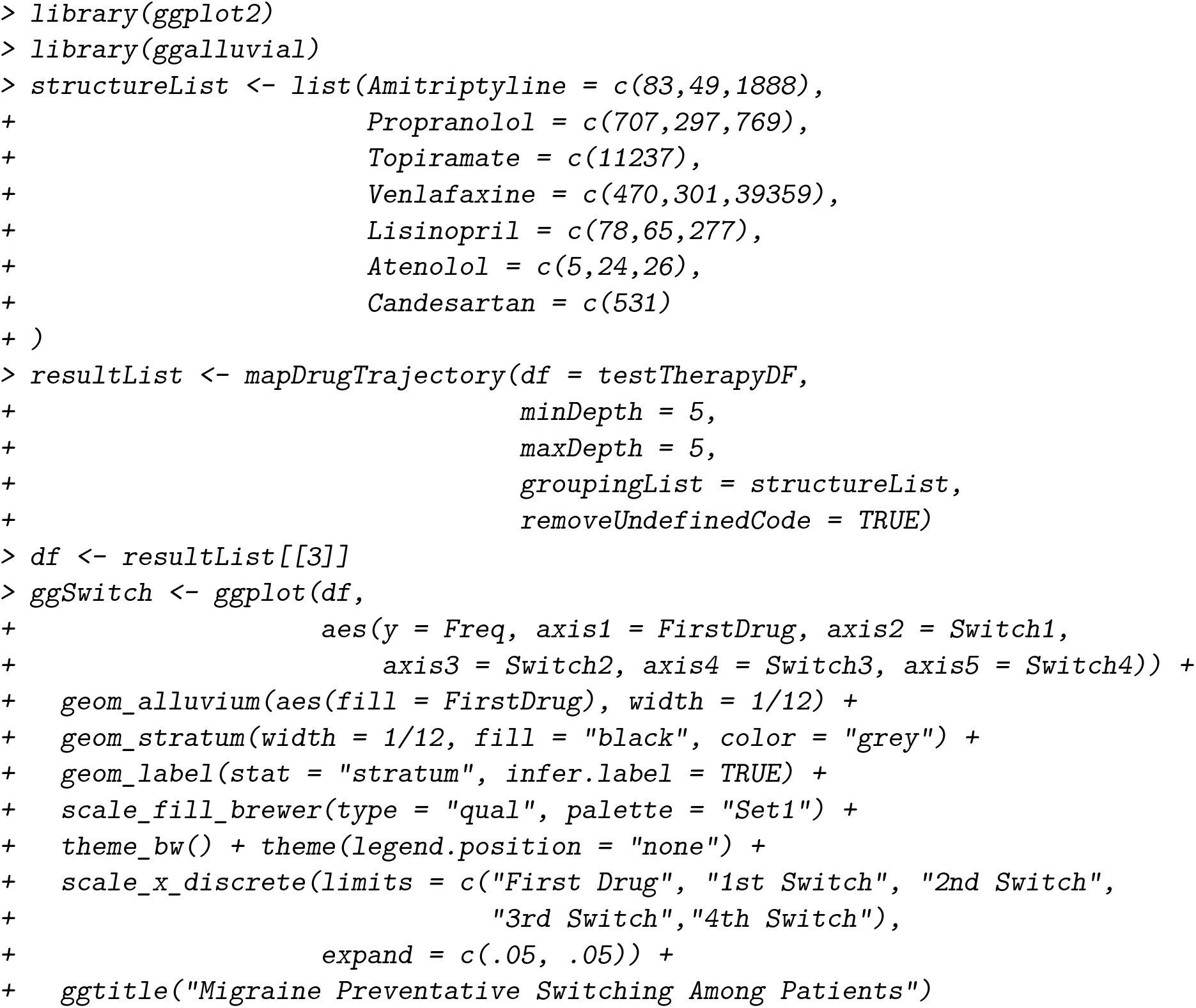

### 3.5. Prescription timeline construction

**rdrugtrajectory** contains several functions that transforms patient data into a format compatible with mean cumulative function (MCF) semi-parametric estimates, prescription persistence, prescription incidence, and survival analysis.

#### generateMCFOneGroup

Prescription events are binned into weekly units to increase the statistical power at each time point. The user presents a group at a time, for example, *all clinical events of male patients with a first-line prescription of amitriptyline for a migraine*. The clinical data has already been refined using the steps for first-line prescription, as described above. The function `generateMCFOneGroup` accepts a dataframe or events, the MCF start date (*eventdates* are adjusted so all patient records in the dataset begin at the same time), and the minimum number of events per patients (by default this is two events). The following example presents the calculation of first prescription events, the assignment of gender and the calculation of MCF of prescription (*therapy* dataframe) burden of amitriptyline and propranolol:

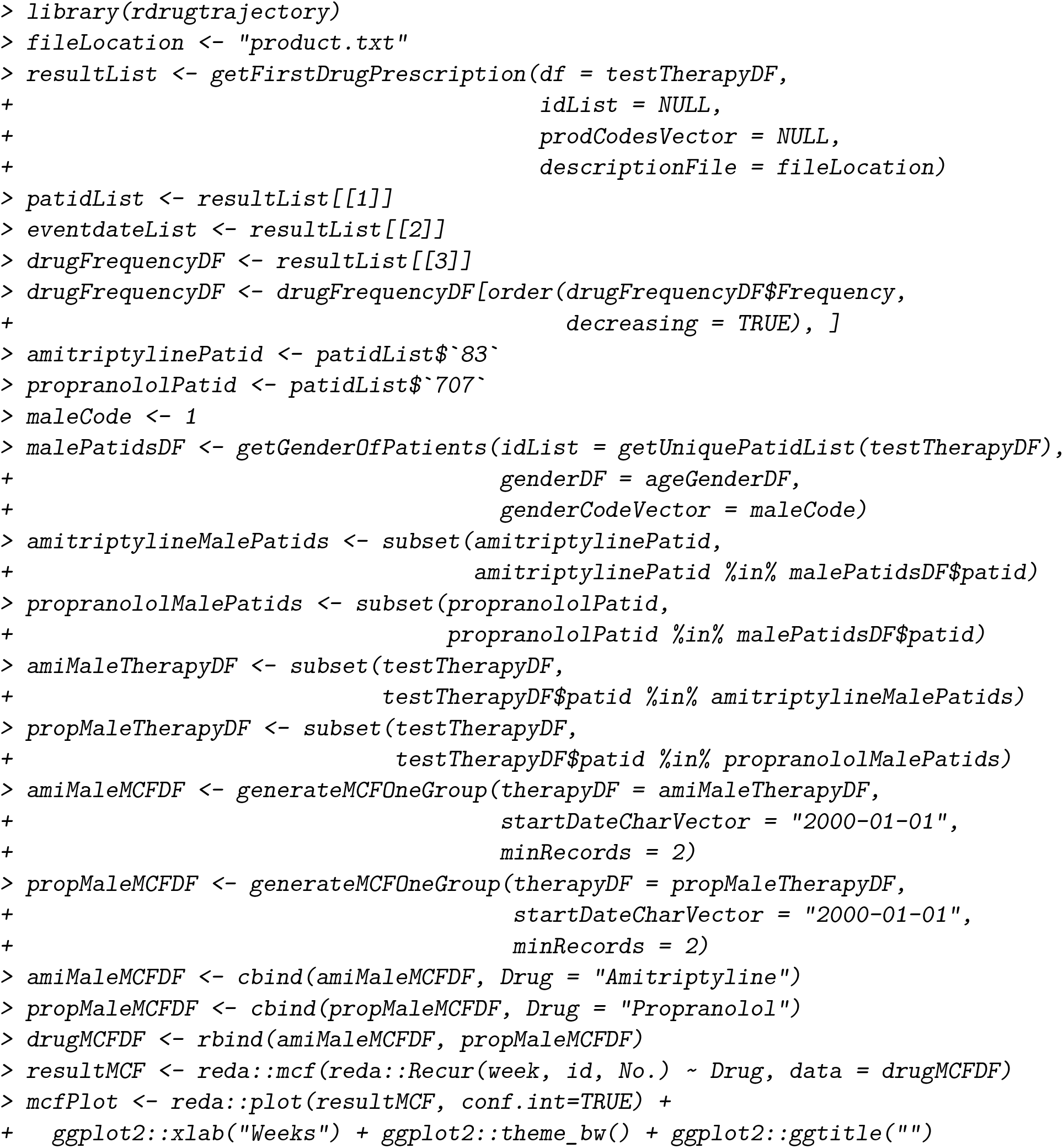

#### getFirstDrugIncidenceRate

Prescription incidence be calculated with `getFirstDrugIncidenceRate`. The following code demonstrates how to use a `FirstDrugObject` to calculate incidence rates for a set of *prodcodes*. The study observation starts from the `enrollmentDate` and ends at the `studyEndDate`:

~~~
*> library(rdrugtrajectory)*
~~~

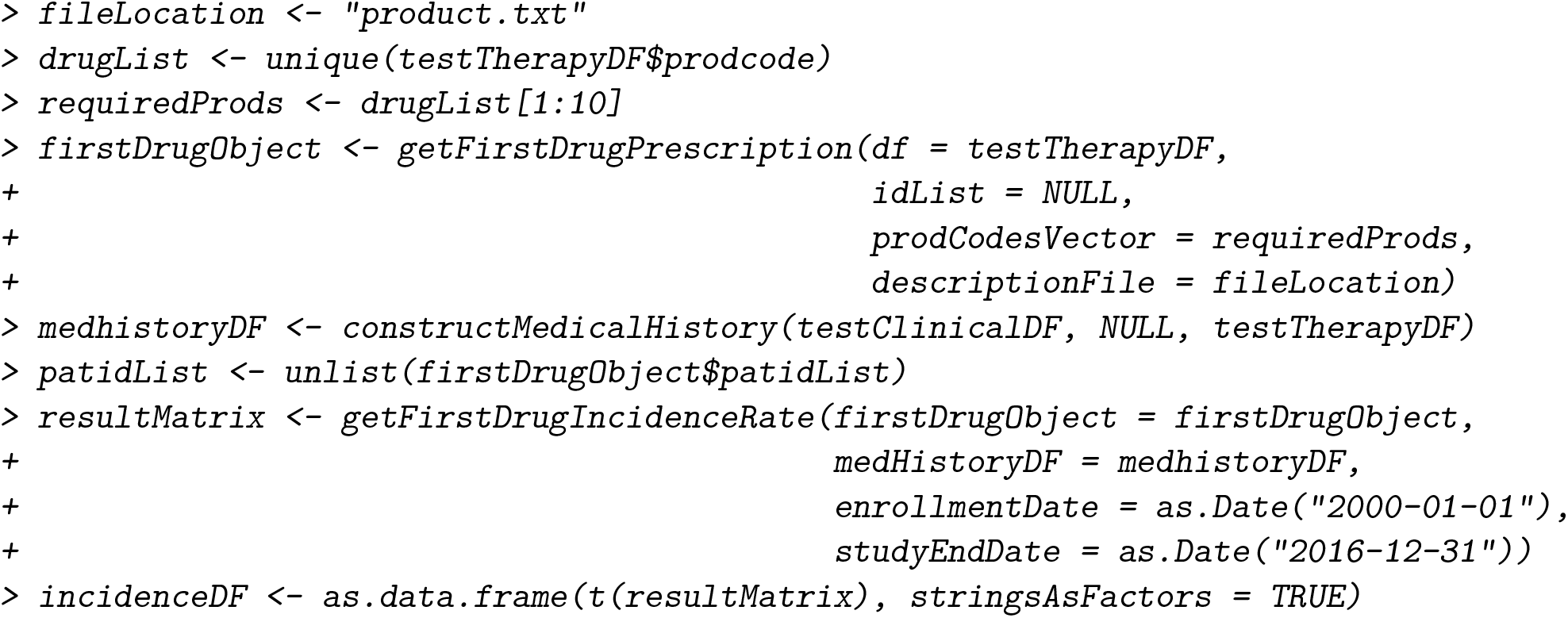

The above example returns an incidence rate of 0.11 per 17 person years over a cohort of 3838 patients. For a detailed description please see *Detail* for `getFirstDrugIncidenceRate` in the reference manual.

#### getDrugPersistence

Prescription persistence is calculated as the fraction of patients with a prescription for a specific treatment *N* -days after the first prescription event. For example, if we wanted to calculate the fraction of patients with a prescription 365-days after their first prescription, with a 30-day buffer either side, one specifies a `duration` of 395-days and a preceding `buffer` of 60-days (therefore, capturing the range 335 to 395, 30-days either side of one calender year):

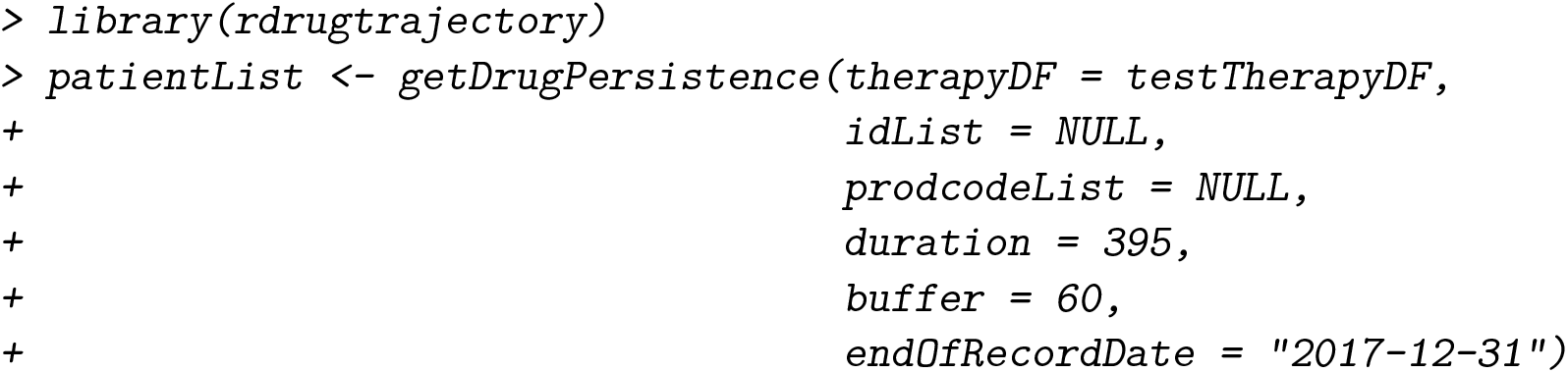

Of 3838 patient *therapy* records, 954 patients had a prescription 365 (+/-30) days after the first prescription event on record, resulting in a crude fraction of only 0.25 patients. `getDrugPersistence` only observes events recorded precisely `duration` days after the first prescription. The `buffer` can be used to identify patients who received a prescription shortly after the end of the `duration`, but more importantly, to ensure patients actively undergoing treatment (indicated by a prescription shortly before the desired `duration` days) are included. As the `buffer` is reduced, the fraction of prescription persistence is reduced until the algorithm attempts to only identify patients with a prescription exactly `duration` of days after the first prescription. Future software updates will incorporate repeat prescription data to increase the accuracy of the calculation.

## 4. Closing remarks and future work

**rdrugtrajectory** is an R package which has the potential for exciting applications such as improving clinical decision-making, identifying possible new treatments and analysing outcomes from existing treatments. We have demonstrated several functions, some of which detail sorting and matching records whilst others demonstrate fundamental statistical analysis. We used fabricated clinical and prescription dataframes, along with the age, gender and index of multiple deprivation score of each patient and presented analyses of cohort-wide prescription patterns, first-line treatment distributions, how to stratify by patient characteristics, and some basic tools to assist longitudinal analysis of prescriptions.

The descriptions presented in this publication are not substitutes for the material in the reference manual. We recommend the reader consults the R `?` help command or reference manual before running a function. In particular, functions related to the construction of timelines for survival analysis (time dependent/independent Cox regression, Kaplan Meier survival curves and mean cumulative function) or a matrix for drug incidence rate requires fine tuning of several parameters.

The latest release of **rdrugtrajectory** along with source code and reference manual is available for download from https://github.com/acnash/rdrugtrajectory. Whilst active members of the scientific research community we will continue to add new features to **rdrugtrajectory** whilst making necessary improvements to existing features.

## Acknowledgements

Oxford Science Innovation, NIHR Oxford Biomedical Research Centre and NIHR Oxford Health Biomedical Research Centre (Informatics and Digital Health theme, grant BRC-1215-20005). Thanks to Dr Michelle Hardy for assistance with the article.

